# Early cell autonomous and niche-mediated alveolar epithelial response to influenza infection in primary lung organoids

**DOI:** 10.1101/2025.10.24.684481

**Authors:** Amber Elitz, Sharlene Fernandes, Kathleen C.S. Cook, Helen I. Warheit-Niemi, Barbara Zhao, Andrea Toth, Amanda L. Zacharias, William J. Zacharias

## Abstract

Influenza A virus (IAV) infection is a significant cause of morbidity and mortality for patients worldwide. Alveolar type 2 (AT2) cells are the preferential target of IAV as part of the pathogenesis of viral pneumonia and acute respiratory distress syndrome (ARDS). Early IAV infection of alveolar cells has been challenging to model both *in vitro* and *in vivo*. To address this challenge, we used a combination of murine and human primary alveolar organoids to define methods for robust IAV infection and evaluated cell-autonomous consequences of IAV using a temporal series of multiome paired single nuclei RNA/ATAC sequencing. Infected AT2 cells undergo conserved changes defined by early loss of surfactant secretion, decreased lipid biogenesis, a rapid burst of antiviral response, and late viral-mediated suppression. Surprisingly, uninfected AT2 cells undergo substantial transcriptional and epigenomic changes in IAV-treated cultures, leading to transition to damage-associated cell states within hours via a process driven by the inflammatory milieu of murine organoids. Together, these data provide new methods for high-fidelity modeling of IAV infection in alveolar cells and define a conserved AT2 cell response signature to IAV with implications for ARDS pathogenesis.

## Introduction

Influenza A viruses (IAV) infect nearly a billion people annually, and over 3 million of those infections cause severe lower respiratory tract disease, or viral pneumonia^1,2^. Viral pneumonia leads to disruption of the critical alveolar gas exchange surface via direct infection of alveolar type 2 pneumocytes (AT2) in the distal lung. Many viral pneumonia patients will develop respiratory failure and require ICU care due to acute respiratory distress syndrome (ARDS)^3^. ARDS is characterized clinically by bilateral pulmonary infiltrates and hypoxemia resulting from fluid and immune cell influx into the alveolar space and associated alveolar collapse (atelectasis)^4,5^. Coordinated regeneration is required to restore gas exchange and alveolar structure^6^. Failure of these regenerative mechanisms is associated with increased risk of secondary bacterial pneumonia^2^ and pulmonary fibrotic remodeling leading to fibroproliferative ARDS^7^. These complications are associated with high morbidity and mortality.

Previous studies have attributed the extent of injury following IAV infection and risk of subsequent ARDS to many factors^2^, including immune-mediated tissue damage, lysis of infected cells impairing the alveolar barrier, loss of vascular integrity, and physiological stress resulting from hospital interventions like mechanical ventilation^4,5,7-9^. The knowledge that these factors contribute to disease has not yet led to improved anti-influenza therapies. Extensive literature has highlighted that even patients who recover from viral pneumonia may have significant, long-lasting respiratory dysfunction^7,10,11^. These limitations suggest that new approaches are needed to study IAV disease pathogenesis prior to onset of ARDS, and to guide the development of new therapies.

Pathologically, IAV causes extensive injury to the alveolar epithelium during viral pneumonia. The primary role of the alveolus is to maintain ventilation via the air-facing gas exchange surface^12^, a role which requires high surface area alveolar type 1 (AT1) cells to interact with alveolar capillaries to provide functional ventilation. AT2 cells produce surfactant proteins to regulate surface tension in the alveolus, preventing atelectasis^13^. These cell populations are maintained through activity of Wnt signaling in AT2 progenitors^14,15^, which can enter the cell cycle, replicate, and replenish AT1 and AT2 cells following IAV injury of the distal lung. After infection, the IAV genome replicates in the nucleus of infected cells. Recent studies have shown that host stress^16^ and viral Nonstructural Protein 1 (NS1) contribute to readthrough error and repression of host transcription in infected cells, leading to changes in cellular chromatin structure and gene expression^17,18^. AT2 cells additionally play a vital role in pulmonary immunity^19^. During IAV infection, virally infected cells present IAV-derived antigens at the cell surface, activating innate immune cells in the alveolus and leading to recruitment of circulating immune cells which target and clear infected cells. Both tissue resident and recruited immune cells secrete cytokines which signal to other cells in the infected alveolus^20^. Thus, damage to alveolar epithelial populations, either through direct, cell-autonomous mechanisms or the inflammatory milieu, results in the loss of alveolar gas exchange, regenerative function, or surfactant secretion. These cellular processes underlie key aspects of the pathogenesis of ARDS^21^. However, the direct alveolar epithelial-specific dynamics of IAV infection are challenging to study *in vivo* due to immune-mediated injury and clearance of infected cells. Conversely, existing two-dimensional *in vitro* systems do not model the cellular relationships of the alveolar epithelium, require viral activation to improve infectivity of cells, and poorly model AT2 cell state and function.

To address these challenges, we present here findings describing optimization and modeling of IAV infection in primary murine and human alveolar epithelial organoids. Using this *in vitro* infection platform, we characterized both cell autonomous and paracrine effects on cell state, transcriptome, and epigenome using multiomic single nuclei RNA and ATAC-seq. We find that IAV-infected AT2 cells differ transcriptionally and epigenomically from uninfected AT2 cells, and that many of these changes are driven by inflammatory signaling occurring within 24h of IAV exposure *in vitro*. Using paired analysis of RNA expression and chromatin accessibility data, we define a conserved AT2 response to infection associated with changes in surfactant production and immune-mediated anti-viral responses, delineating key IAV-mediated mechanisms underlying ARDS pathology and providing a large, integrated multiomic dataset of IAV-mediated for exploration and community use.

## Results

### Optimizing influenza infection in complex lung alveolar organoids

To model IAV infection in alveolar epithelial organoids (AEOs), we generated AEOs by sorting primary AT2 progenitor cells from adult mice and combining with expanded primary mesenchyme in Matrigel and SAGM as previously described^22^. After culturing AEOs to 21-28 days, we attempted infection in organoids still within the Matrigel plug by adding H1N1 PR8 IAV at an multiplicity of infection (MOI) of 10 (1.025×10^6^ PFU) to the top of each well in spiked Small Airway Growth Media (SAGM). We observed that few cells were infected and expressing IAV nucleoprotein (NP) 24 hours post infection (hpi) (Figure 1A), suggesting that viral diffusion through the Matrigel was too inefficient for robust AEO cell infection. To improve infection kinetics, we next removed the mature organoids from matrix and transitioned the isolated organoids to suspension culture in SAGM. We evaluated protein expression of AGER and surfactant protein C (SPC) to detect AT1 and AT2 cells, respectively. Transition to suspension minimally disrupted organoid cell number and cavity integrity within the first 24h, with a general loss of AT1 cells and a near significant loss of cell number per organoid by 36h (Figure 1B-C). We therefore chose to restrict our evaluation to the first 24h of suspension culture. This had the advantage of focusing our analysis on early impact of IAV.

**Figure 1.**
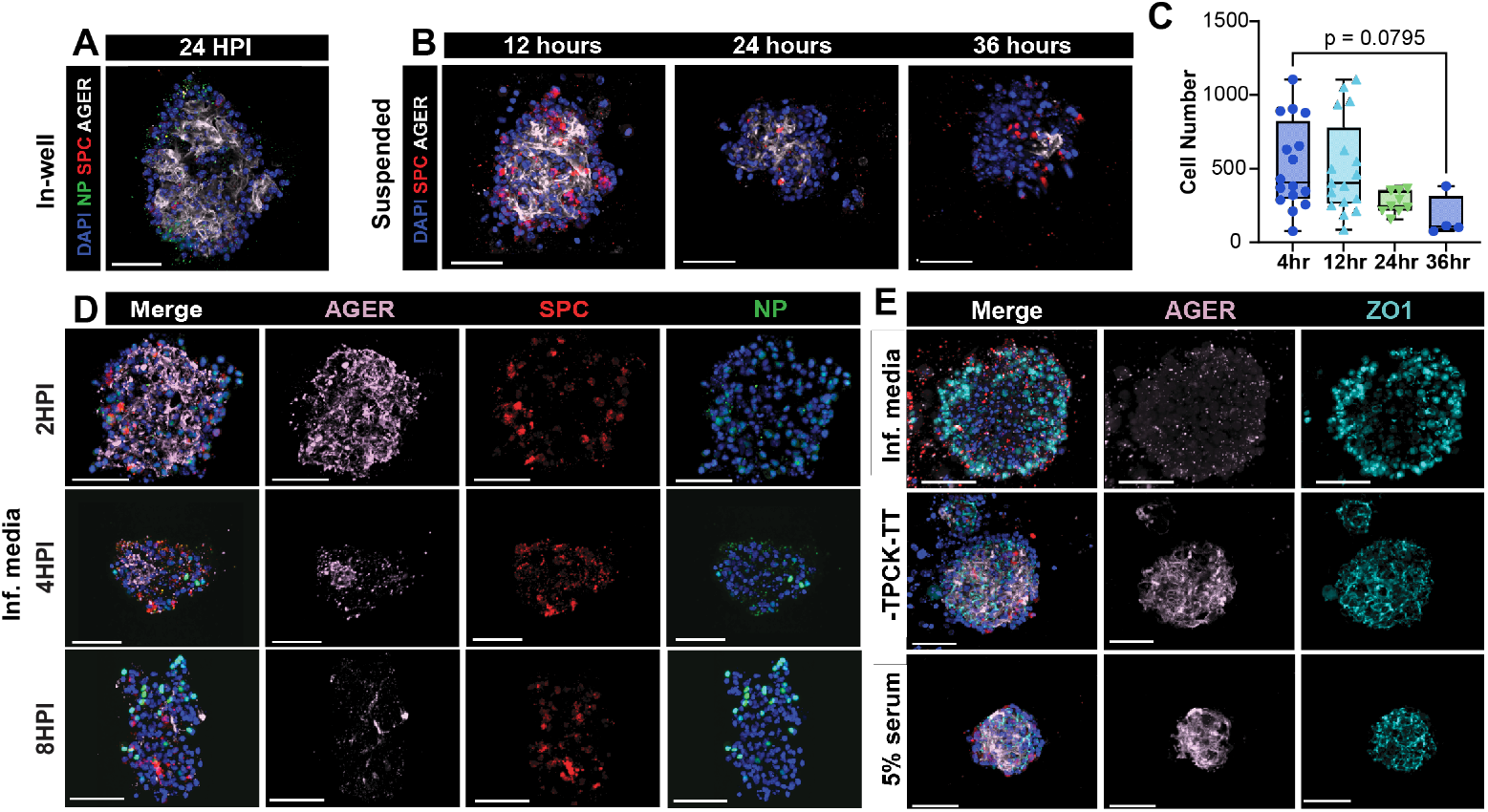
Optimization of Influenza A (IAV) Infection Conditions for Alveolar Organoids (AEOs). (A) Treatment of AEOs with IAV in media placed on apical surface of transwell above matrigel plug. Minimal NP expression is evident at 24hpi. (B) Morphology of AEOs in suspension culture after liberation from Matrigel. AT1 and AT2 cell morphology is maintained for 24h post-suspension but has decreased by 36h. (C). Cell number per organoid in suspension culture. We noted a nearly significant loss of cell number by 36h post liberation. (D) Cell state of AEOs with infection of IAV using SAGM base media with TPCK-treated trypsin and lacking serum (Inf. media). Suspension in this medium allowed flu infection by 8h (NP) but led to loss of AT1 cells (AGER) without loss of AT2 cells (SPC). (E) Impact of different media compositions on cell junctions in AEOs. Infection media (top row) led to loss of AGER and associated loss of ZO-1 marked cell surface tight junctions. Removal of TPCK-treated trypsin or addition of 5% serum prevented these changes. Scale bars = 50 μm. *(SPC = Surfactant Protein C [AT2 marker]; Ager = Advance Glycosylation End-product Specific Receptor [AT1 marker]; NP = IAV nucleoprotein [infected cell marker], ZO-1 = Zonula Occludens-1 [cell surfact tight junction marker])*.

We next sought to optimize our AEO infection media conditions. Published protocols in monolayer culture and lung spheroids have demonstrated that TPCK-treated trypsin cleaves influenza entry factor HA0 to expose the viral particle binding sites and facilitates infection initiation in cell lines lacking key transmembrane proteases^23^. The addition of serum to infection media for IAV has been suggested to impede viral infection, likely by quenching trypsin activity^23^. We therefore evaluated an “infection media” that was both serum-free and contained TPCK-treated trypsin. We found that AEOs, particularly the AT1 population, were extremely sensitive to disturbances in mechanical tension induced by trypsin. While the organoids retain AT1 structure and specification in suspension (Figure 1B), the addition of TPCK-treated trypsin cleaved cell-cell junctions and produced a selective loss of AT1s followed by general organoid degradation within 8h (Figure 1D). Removal of trypsin from the media or addition of 5% serum for neutralization protected AT1 morphology in AEOs (Figure 1E). We therefore decided to proceed with infection modeling in suspension culture using complete SAGM.

### Infection dynamics and cellular specificity of IAV in vitro in AEOs

Using our optimized infection conditions, we performed an infection timeseries in AEOs in SAGM suspension (Figure 2A) at 8, 16, and 24hpi, and followed infection kinetics by whole mount IHC to detect (Fig 2B-E) and quantify (Figure 2F-G) NP-label nuclei. We reasoned that by 24hpi, host cells have undergone a full viral life cycle with the majority of infected cells experiencing a period of continuous viral replication^24^, suggesting that key changes in early IAV pathogenesis would likely be captured by this timepoint. The percentage of NP^+^ cells per organoid progressed from <10% at 4hpi to 24% at 8hpi, 29% at 16h, and >40% at 24hpi (Fig 2F). These results suggest that cells expanded in primary AEO culture retained sufficient expression of key transmembrane proteases that perform the HA cleavage to enable viral binding in the absence of TPCK-treated trypsin. IHC demonstrated infection primarily in AT2 cells when corrected for population size (Fig 2G). To assess viral binding availability, we performed staining with the lectin Sambucus Nigra Agglutinin (SNA) to evaluate the presence of the primary IAV binding moiety, α2,6 sialic acid (Fig 2H). AT2 cells demonstrated increased SNA labeling compared to AT1 in AEOs. This was confirmed by by flow cytometry from freshly isolated epithelium from wild type mice (Figure 2I). To confirm this specificity in cell infectivity *in vivo*, infected R26R-EYFP mice with a PR8 strain harboring Cre and evaluated YFP expression per cell type by flow cytometry (Figure S1). We confirmed significantly more infected AT2s than AT1s 72hpi (Fig 2J). Taken together, these data demonstrated a clear progression of IAV infection in AEO across the 24h in suspension, which replicated the cellular targeting of IAV in the lung *in vivo*.

**Figure 2.**
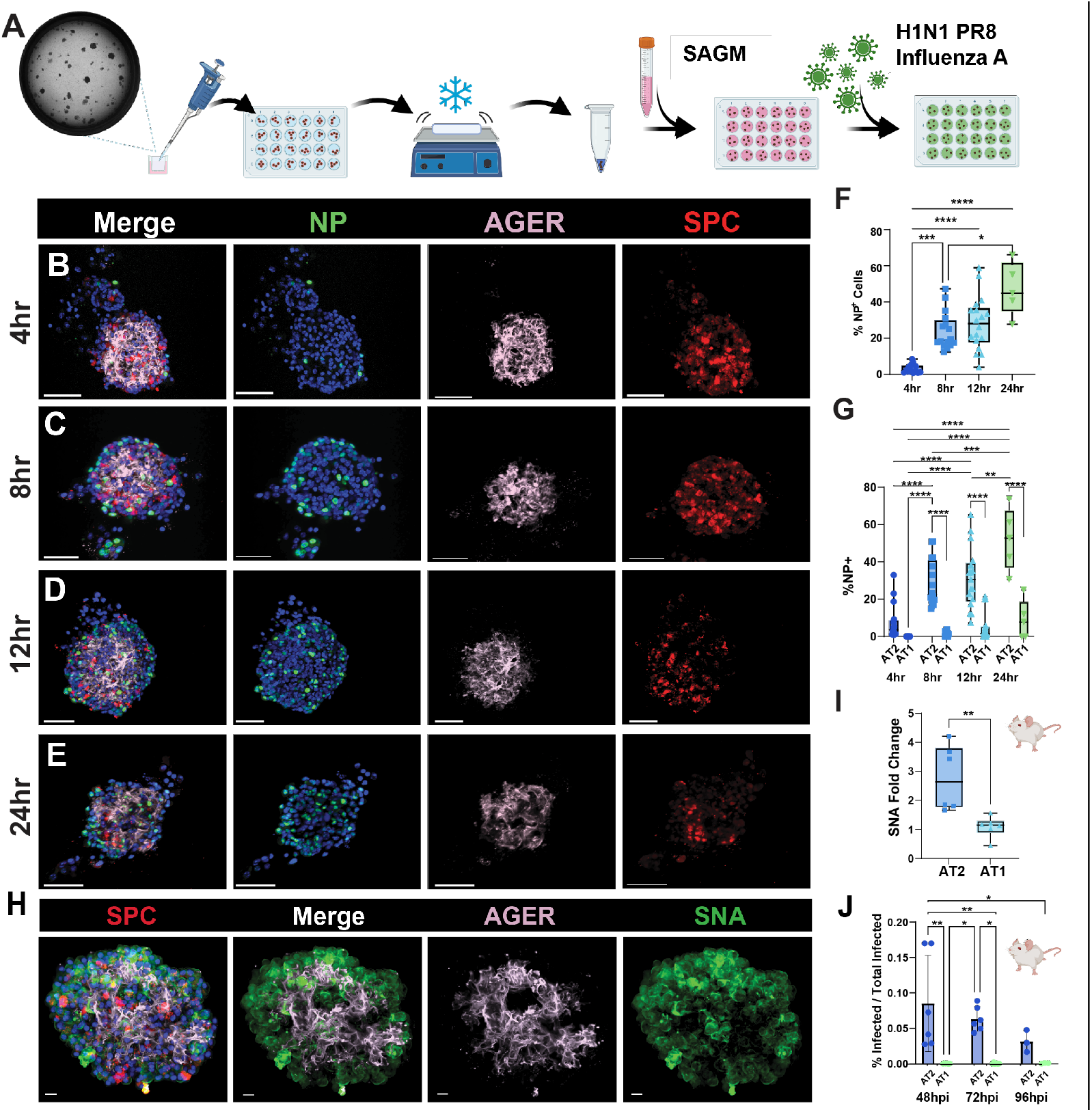
Temporal dynamics and cellular specificity of IAV infection in AEOs. (A) Schematic of method for liberation and treatment of AEOs with IAV. (B-E) AEOs treated with H1N1 develop progressively increased influenza infection as measured by nuclear expression of H1N1 NP across 24hpi. (F) Quantification of NP-positive nuclei at each timepoint of infection. (G) Cell type specific infection in AEOs. Significantly more AT2 are infected than AT1 by 8hpi and thereafter. (H) Expression of the flu binding α-2,6-sialic acid in AEOs marked by expression of sambuccus nigra lectin predominantly in AT2 cells. (I-J). *In vivo* confirmation of SNA expression (I) and increased infection rate (J) in AT2 cells during early flu. Scale bars = 50 μm. *(SPC = Surfactant Protein C [AT2 marker]; Ager = Advance Glycosylation End-product Specific Receptor [AT1 marker]; NP = IAV nucleoprotein [infected cell marker], SNA = Sambuccus nigra lectin [flu binding motif α-2,6-sialic acid])*. P values by ANOVA with multiple comparison (F,G,J) or T test (I): * = <0.05, ** = <0.01, *** = < 0.001, **** = < 0.0001.

### Defining trajectories of cellular injury in IAV AEOs with multiome sequencing

To evaluate changes in gene expression and chromatin accessibility in infected AEOs, we performed multiomic single-nuclei RNA (snRNA) and ATAC (snATAC) sequencing on nuclei isolated from AEOs suspended in SAGM with or without H1N1 at 8hpi, 16hpi, and 24hpi. The multiome assay generated linked RNA and ATAC data at per-nuclei resolution, allowing us to assess both gene transcript expression and chromatin accessibility per cell. To identify infected cells, we generated a custom mm10^25,26^ genome alignment file which included the PR8 viral genome to allow detection of cells with active viral replication occurring in the nucleus. After filtering to remove doublets^27^ and ambient RNA^28^, we recovered 33,926 single-nuclei transcriptomes with a range of 3,642 to 8,793 cells per timepoint and condition with a median of 943 unique molecular identifiers (UMI) (interquartile range 614 – 1,712) per cell, and a median of 734 genes (interquartile range 509 – 1,186) per cell. These RNA sequencing depths were below those previously observed with single-cell RNAseq in AEOs^22^ or in vivo after flu infection^21,29^, but provided resolution to define infected cells and simultaneously compare transcriptome and epigenome state. We focused our analysis on major cell states defined by nearest neighbor analysis using both snRNA and snATAC assays.

Following unbiased clustering (Figure 3A), we identified AT2 cells, as well as a population of AT1 and AT1-like (AT1tr) cells within the epithelium, as expected based on our IHC results (Figure 3B). We also detected the anticipated mesenchymal/feeder cell population, including mouse lung fibroblasts and some myeloid cells that persist in the feeder culture. These cells were identified by marker gene analysis as most similar to interstitial macrophages^30^. Unique to the IAV-exposed condition, we noted the presence of two populations we denoted as stressed transitional epithelium (STE). These populations share a gene expression signature typical of previously published^31^ Krt8^High^ (also known as PATS^32^, DATP^33^, or ADI^34^) cell populations known to arise in diverse forms of lung epithelial cell injury. The presence of these cell states has been described within several days of injury *in vivo*, but they arise within hours following IAV infection in our AEO system. Further, we detect a distal secretory-like population defined by joint expression of Sftpc and Scgb1a1, most prevalent at the 24-hour timepoint in unexposed AEO (Figure 3A-B). This population appears to arise with acquisition of Scgb1a1 expression in AT2 during the unexposed suspension, potentially as a response to suspension culture in the absence of other stressors (Figure 3C). We note that there is likely a higher degree of heterogeneity of cell states present within both the epithelial and fibroblast populations, but the depth of multiome nuclear RNA sequencing was insufficient for confident definition of these more refined cell states.

**Figure 3.**
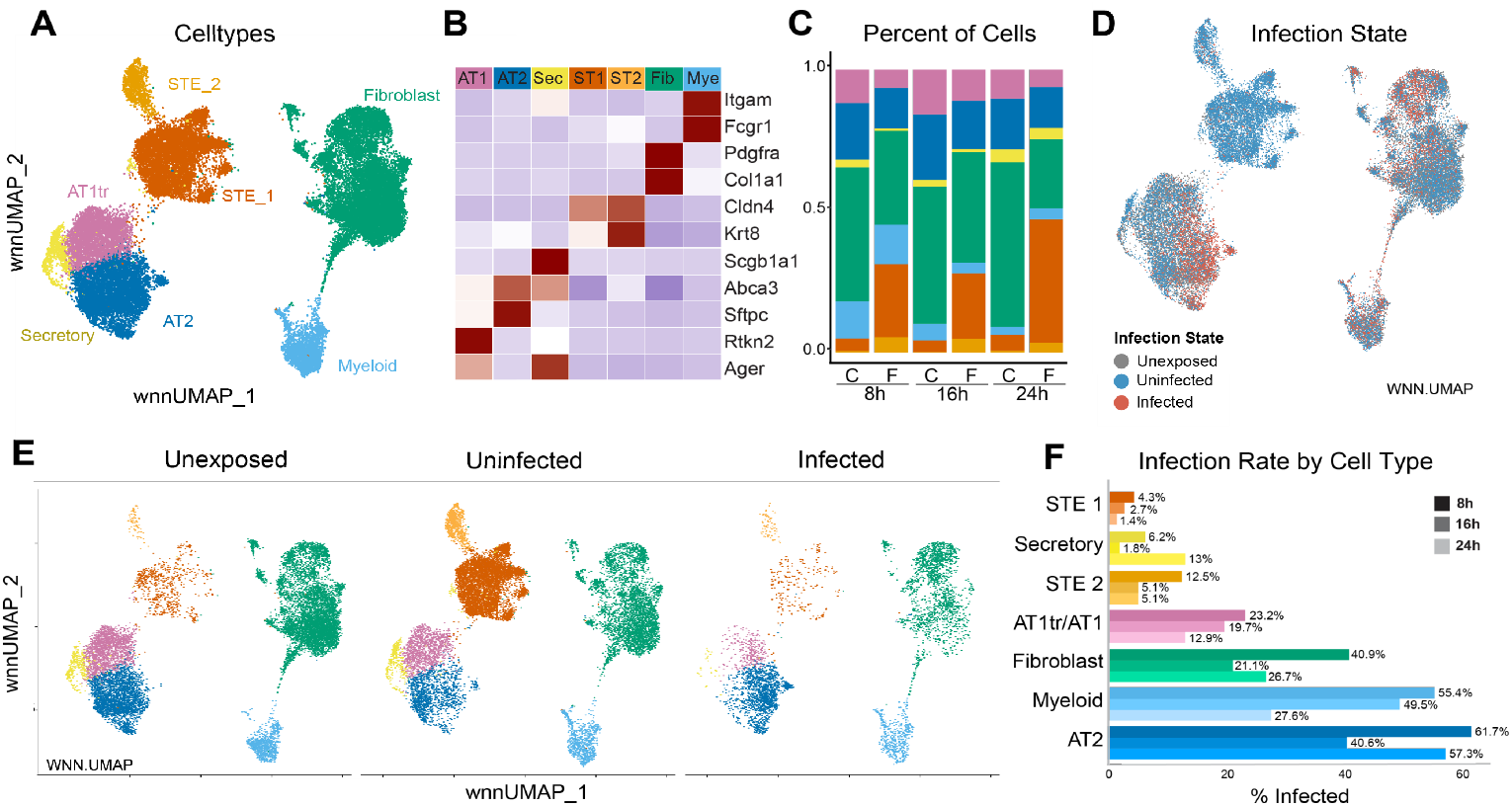
Single nuclear analysis of flu infection in AEOs. (A) UMAP of cells from unexposed and flu-treated AEOs. (B) Cell state markers defining cell states shown in (A). (C) Frequency of cell states observed in each unexposed [C] and flu-treated [F] AEOs. (D-F) Infection states. Overall UMAP (D), individual condition UMAPs showing unexposed AEOs, uninfected cells in flu-treated AEOs, and infected cells in flu-treated AEOs (E), and overall rate of infection per cell type (F). *AT1tr = AT1-like/transitional cells, STE = stressed transitional epithelial cell. Other abbreviations per main text*.

Next, we identified the IAV infection state in cells. We generated an IAV expression score based on the number of IAV genes and expression levels to distinguish three classes of cells (see Methods): 1) “Unexposed” (UE) cells from IAV-naive AEO cultures, 2) “Infected” cells which had high levels of flu-specific gene expression and 3) “Uninfected” (UI) cells which were exposed cells in IAV-treated cultures but did not meet the IAV gene expression thresholds for infection (Figure 3D-E). Concordant with our organoid and *in vivo* IHC data, the majority of infected epithelium were AT2 cells with few infected AT1 cells. Surprisingly, we noted the presence of substantial populations of infected fibroblasts and myeloid cells (Figure 3E) with a highest enrichment at the 8h timepoint (Figure 3F). We suspect that these “off-target” infections are due to prolonged saturation with viral particles with no protective extracellular matrix. HA cleavage by epithelial cells may also improve viral availability for infection of mesenchymal lineages. Notably, both STE populations had a very low rate of flu infection.

### Gene expression profiles of infected cells within AEO

We next turned our attention to defining the pathways associated with infection state in AT2 cells. Within the ‘Infected’ population, we defined cells as “Early” infection based on early gene signatures (expressing PB1 and 2, PA, and PA-X, NP, NS1, and NEP) for genes involved in poly-A cap snatching, host machinery hijacking and disruption, and nuclear transport. We scored cells as “Late” infection due to expression of later stage genes (M1 and M2, HA, NA, and PB-1-F2) involved in building the viral particle membrane and membrane proteins in addition to the ‘early’ genes (Figure 4A-C)^2^. The resulting patterns of infection stage at each timepoint aligned with timelines previously reported in monolayer cell culture models^24^. In AT2 cells, we detected ‘Early Infected’ cells at each timepoint. We interpreted these results to indicate that cells experience ongoing infection while in suspension in IAV-media. We detected ‘Late Infected’ cells 8hpi, decreased at 16hpi, and then again increased at 24hpi; these data suggest that our system allowed up to two full cycles of viral replication by 24hpi (Figure 4B-C).

**Figure 4.**
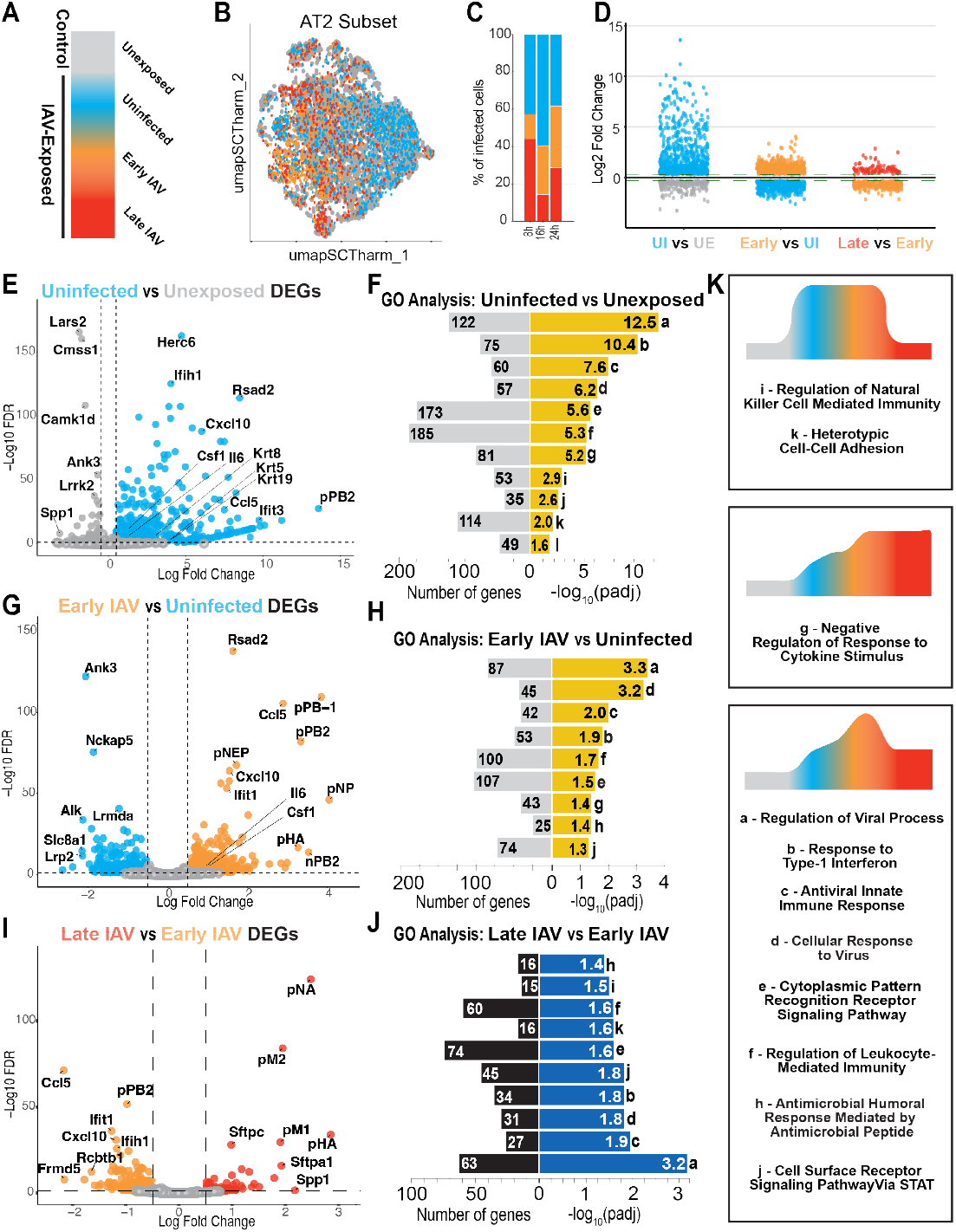
Trajectory of gene expression during IAV infection in AEOs. (A) Schematic color scheme of cell states in AEOs. Unexposed = grey, Uninfected = blue, Early IAV = orange, Late IAV = red. (B-C) UMAP (B) and frequency (C) showing infection states of AT2 cell subset of AEOs. (D) Strip plots showing number of differentially expressed genes (DEGs) and degree of enrichment across AT2 cell states. (E-F) Volcano plot of DEGs (E) and GO Biological Process enrichment (F) of genes in Unexposed (grey) vs Uninfected (blue) AT2. (G-H) Volcano plot of DEGs (G) and GO Biological Process enrichment (H) of genes Uninfected (blue) vs Early Infected (orange) AT2. (I-J) Volcano plot of DEGs (I) and GO Biological Process enrichment (J) of genes in Early Infected (orange) vs Late Infected (red) AT2. Upregulated processes are shown in yellow, and downregulated processes in blue. (K) State dynamics of enrichment of GO terms from F, H, and J. Letter to left of process name matches letters to right of bars. Some terms are enriched at multiple timepoints. Diagrams in K show dynamics of change of associated terms (boxes) across cell states in AEOs.

To define the cell-state specific effects of IAV infection on AT2 cells, we created pseudobulk profiles of cells scored as Unexposed, Uninfected, Early IAV, or Late IAV infection, ensuring even contribution of cells from each timepoint into the artificial replicates to prevent timepoint biases. We identified differentially expressed genes (DEGs) which we defined by an adjusted p value of <0.05 and a fold change of >1.25 (Figure 4D) within these populations. Surprisingly, we noted the most DEGs comparing the Unexposed to Uninfected cells, with fewer changes comparing Uninfected to Early Infection, and limited significant genes changed between Early and Late infection.

To evaluate major pathways associated with genes regulated across the course of AT2 infection, we performed Gene Ontology (GO) Biological Process (BP) term enrichment analysis^35,36^. Uninfected AT2 cells upregulated genes associated with pathways of AT2 stress response and immune signaling, including viral and interferon response (Figure 4C-D). These terms were driven by leading edge genes including Krt8, Krt19, Cxcl10 (IP-10) and Ccl5 (RANTES) (Figure 4E and F). Many of these genes and pathways were further upregulated in Early IAV cells compared to Uninfected AT2 (Figure 4G-H), but then downregulated in Late IAV (Figure 4I-J). The Late infection condition was instead defined by genes associated with pathways controlling protein folding, with higher expression of several heat shock protein genes and ER stress markers (Figure 4I,K). Finally, we noted a significant initial increase in intracellular markers of type 1 interferon response in AT2, including Irf5 and Isg15. Consistent with published results, this signature is then downregulated in late infection, which has been reported as a consequence of viral-mediated suppression^37^. Taken together, these data unite several proposed mechanisms underlying infleunza response in AT2 cells^37-40^ (Figure 4K).

### Cell epigenomic state changes in IAV infection

We next turned our attention to the epigenetic consequences of IAV infection by comparing Unexposed, Uninfected, and Infected AT2 cells in snATACseq data (Figure 5A). We identified differentially open regions of chromatin with a logFC >0.25 and an FDR threshold of 0.05 within pseudobulk chromatin signatures from each of these cell states. Using an ArchR^39^-like approachto identify the top 25% most active differentially regulated peaks (Figure 5B), we the largest difference in open chromatin between Uninfected versus Unexposed AT2 (Fig 5C). Taken with the snRNA findings, these data implied that a majority of chromatin regulation was related to detection and early response to IAV in the system rather than direct infection in AT2 cells. We found significant changes in chromatin near MHC- and IRF-related genes in Uninfected cells. These genes are important for AT2 immune function and the associated RNAs are significantly upregulated in Early Infected cells (Figure 5C and 4G). GO Terms associated with these differential peaks included terms associated with response to IL1β, TNFα, and suppression of viral genome replication. Chromatin in Uninfected AT2s was also more open near genes associated with keratinization, intermediate filament process, cellular response to inflammatory stimuli, and antigen presentation concordant with the observed significant changes in gene expression by snRNAseq (Figure 5D and 4C).

**Figure 5.**
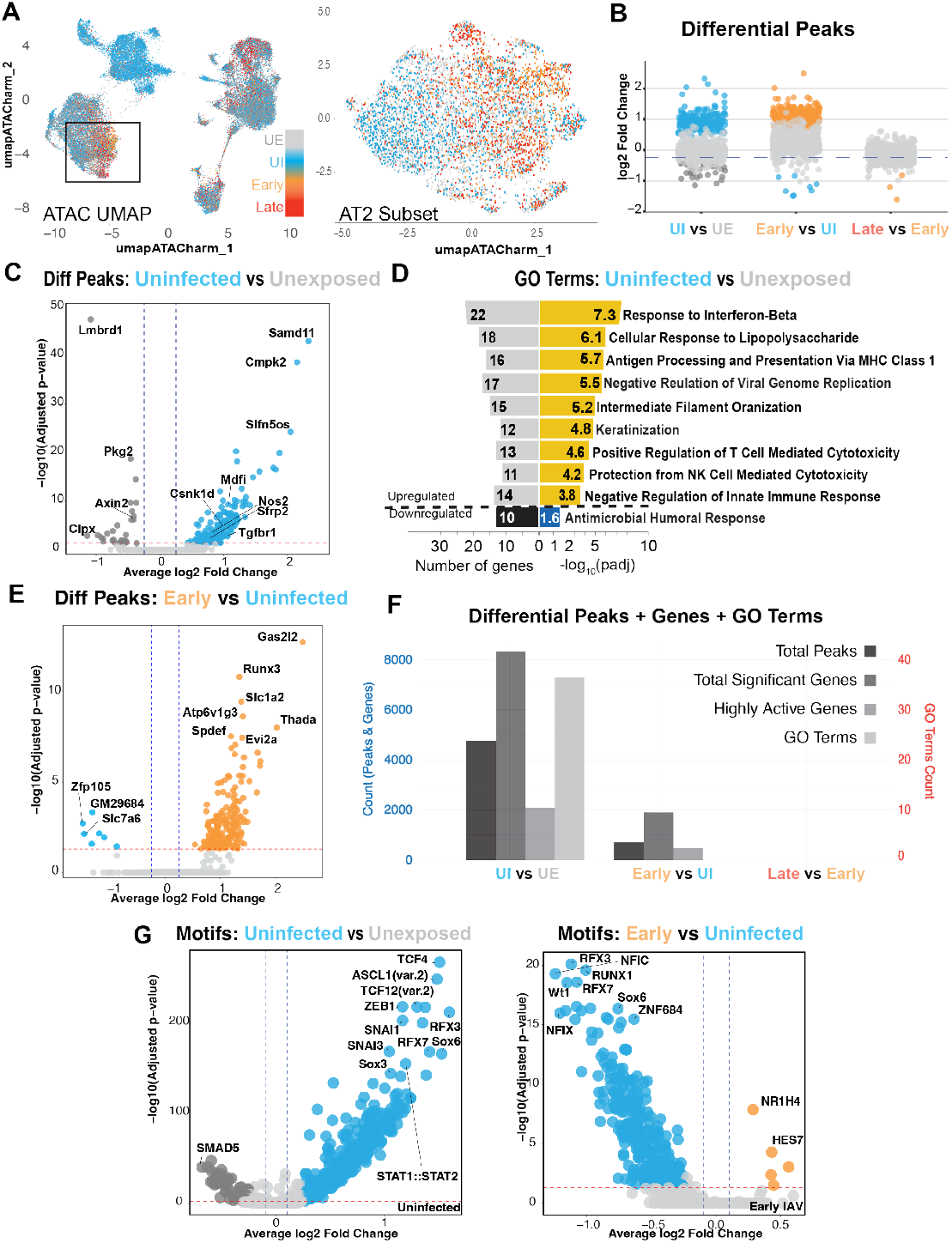
Dynamics of chromatin changes during IAV infection in AEOs. (A) Overall UMAP and AT2 only UMAP based on ATAC assay showing cell infection state. (B) Strip plot showing number and significance of differential peaks across AT2 cell states. (C-D) Volcano plot (C) and GO Terms (D) showing nearest neighbor genes to differentially open chromatin enriched in Uninfected compared to Unexposed AT2 cells. (E) Volcano plot of nearest neighbor genes to differentially open chromatin in Early infected compared to Uninfected AT2 cells. (F) Bar chart showing the number of differentially open peaks, nearest neighbor genes, and associated enriched GO Terms. No enriched GO Terms are associated with newly open chromatin in IAV-infected AT2 cells, and no differentially open chromatin regions passed thresholding for Early vs Late infection. (G) Motifs enriched in differentially open chromatin in Uninfected vs Unexposed (left) or Early infected vs Uninfected (right) AT2 cells.

We next turned our attention to chromatin changes induced by flu infection, comparing Uninfected and Early infected cells. We noted a significant number of differentially accessible peaks in Early IAV (Figure 5E), but noted that these newly accessible peaks were not associated with enrichment of any specific GO BP terms (Figure 5F). We noted that while some chromatin regions were differentially accessible in Early IAV compared to Uninfected, none of these peaks passed our stringent thresholds for significance (Figure 5F). Recent reports have demonstrated that IAV inhibits transcriptional termination in cells lines, leading to widespread readthrough transcription and significant chromatin accessibility changes^17,18^. While we can not directly assess readthrough transcription in our snRNAseq data, the randomly distributed chromatin changes present in IAV-infected cells likely represent the sequellae of readthrough transcription in infected AT2 cells.

Motif analysis of Unexposed compared to Uninfected AT2s predicted increased activity of TF classes associated with cell shape change (Zeb1)^41^ and MHC class II expression (Rfx3/7)^42^, concordant with the anticipated need for injury response (Figure 5G). These data suggested that following viral detection in the milieu, AT2 cells are primed via inflammatory signaling to increase chromatin accessibility in key immune-activating factors that increase in transcription prior to and immediately following infection. Consistent with the transcriptome and peak accessibility data, motif analysis in Early IAV demonstrated very few enriched binding motifs within these differentially open regions (Figure 5G).

### Stressed Transitional Cells Arise from Uninfected AT2 during response to IAV in AEOs

As noted above, we observed a large population of stressed transitional epithelial cells (STE) exclusively in IAV-exposed AEOs. The STE state has been extensively described in recent pulmonary regeneration literature, where STEs have been shown to both participate in productive regeneration as well as accumulate in more chronic injury^31-34,43-45^. STE arise within days *in vivo*, but were present by 8hpi in AEOs, providing an opportunity to evaluate signatures and possible origins of STEs in IAV-infected AEOs.

Both STE populations were predicted by pseudotime analysis of snRNAseq data to arise from the AT2 population (Figure 6A). Although our system does not include a lineage tracing marker to validate this predicted relationship, uninfected AT2s showed enrichment at the ATAC level for gene programs associated with the transition to STE state, including keratinization and intermediate filament-associated gene categories relative to unexposed AT2s (Figure 5C-D). STE populations in AEOs were defined by RNA expression of cytokeratins 5, 8, and 19 among other established epithelial stress markers (Figure 6B). Both populations expressed components of the “typical” DATP/PATS/ADI/ABI cell^31-34,43-45^ profile described in the literature, however STE2 cells were particularly Krt8^high^ and marked by expression of Cdkn1a, Cdkn2a, and Sfn, markers that have been reported in more pathogenic STE. These cells expressed low levels of the recovery markers Isg15 and Oas1a (Figure 6B), similar to uninfected AT2 cells, suggesting STE do not arise from AT2 cells recovered from IAV infection.

**Figure 6.**
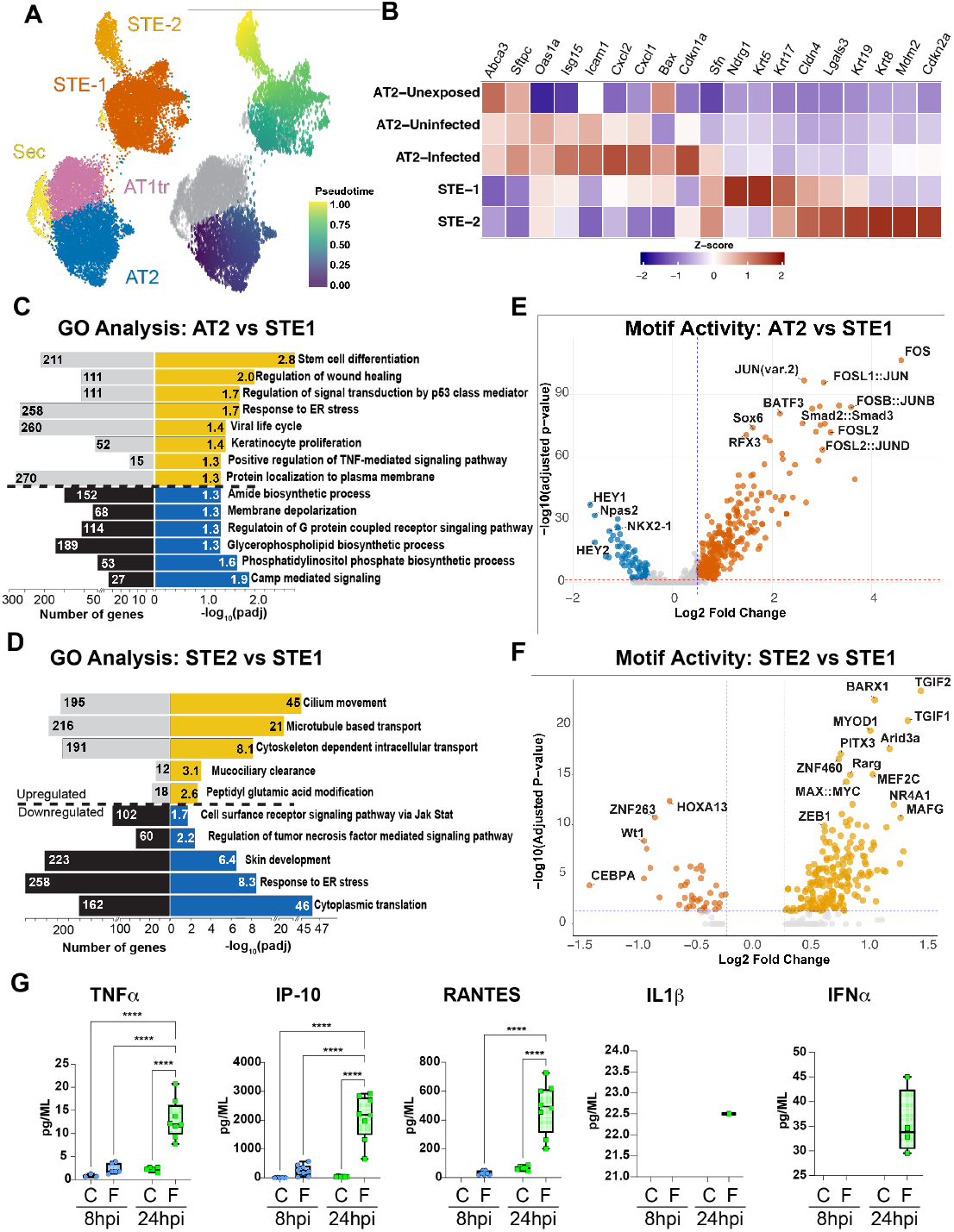
Stressed transitional cells arise from uninfected AT2 driven by inflammatory signaling. (A) Epithelial-only UMAP (left) and psuedotime trajectory (right) suggesting that STE arise from AT2 in IAV-treated AEOs. (B) Marker gene expression of a series of AT2 and STE markers defining cell state in AEOs. (C-D) GO Biological Process Terms based on DEGs for (C) Uninfected AT2s vs STE1s and (D) STE1s vs STE2s. (E-F) Motif activity enrichment based on differentially open chromatin (E) in Uninfected AT2s vs STE1s and (F) and Motif activity enrichment based on differentially open chromatin (F) in STE2 vs STE1. Major reported regulators of STE transition including reduction of Nkx2-1 activity, IL1b signaling driving FOS:JUN signaling, and TGFb signaling are identified in STE1 vs AT2. (G) Cytokine ELISA demonstrating enrichment of multiple IAV-associated cytokines in the media of IAV-infected AEOs. Compare to Figure S4.

When we compared DEGs in STE compared to uninfected AT2s (Figure 6C), we noted enrichment of genes associated with ER stress response pathways, inflammatory signaling, pathways associated with organization of keratins, and p53 signaling. All of these signaling pathways have been associated with STE acquisition in previous literature, supporting the model that STE arise from uninfected AT2s. We noted downregulation of AT2-associated pathways, including lipid metabolism, secretory regulation, and ion transport as well as terms associated with key aspects of ARDS pathogenesis, including local surfactant deficiency and alveolar edema. The shift to STE2 from STE1 was primarily characterized by cytoskeletal, microtubule, and ciliary terms, and a reduction in niche signaling response in the TNF and JAK/STAT pathways (Figure 6D).

### An inflammatory signaling milieu drives STE acquisition in IAV-infected AEO

We next assessed differential motifs found in chromatin open in STE1 relative to uninfected AT2 cells to predict TF activity (Figure 6E). Motif analysis demonstrated a reduction in predicted NKX2-1 activity and increases in predicted regulation by FOS::JUN and SMAD2::SMAD3 motifs in STE1. STE acquisition has been reported following TGFβ signaling upstream of SMAD2/3^31^, and after loss of function of NKX2-1^22^, suggesting similarities in the gene programs driving STE acquisition following IAV-infection in AEOs and in other injury models. Interestingly, STE2-enriched motifs included high predicted activity for the TGFβ target genes and feedback inhibitors TGIF1 and TGIF2 (Figure 6F), suggesting modulation of TGFβ signaling may underlie some differences separating the STE1 vs STE2 states. RNA expression of Tgfb1 was not changed in snRNAseq data and activated Tgfβ ligand was not change by ELISA in stroma exposed to IAV (Figure S2), suggesting that changes in Tgfβ processing or response might be responsible for these changes.

Although the specific trigger for STE acquisition in lung injury is unknown, multiple studies have suggested an inflammation-driven process. Prior literature has directly demonstrated a role for IL-1β^33^signaling in STE transition. FOS::JUN activation is a hallmark of cellular response to IL1B and other proinflammatory cytokines, leading to the hypothesis that inflammatory cytokines induced by influenza may participate in AT2 to STE transitions. To determine which proinflammatory factors are present in the milieu of infected AEO, we compared RNA expression (Figure S3) to protein abundance by ELISA (Figure 6G, Figure S3) for a panel of inflammatory cytokines in IAV treated or untreated cultures. Protein expression of IFNα and IL-1β were only detectable in IAV-treated cultures at 24hpi, and RANTES, TNFa, and IP-10 were detectable at multiple timepoints but significantly upregulated only at 24hpi (Figure 6G). We observed significant upregulation of RNA for multiple pro-inflammatory cytokines in fibroblasts in IAV-infected cultures including the Type 1 interferons IFNα and IFNβ, IL-6, IP-10/CXCL10, and RANTES/CCL5. We also noted increased RNA expression for several overlapping and additional cytokines from myeloid cells in AEOs, including IL-1β, TNFα, RANTES, and MIP1α and MIP1β. These data support the conclusion that the AT2-STE transition in IAV-exposed AEOs was likely driven by inflammatory signaling milieu.

### Generation and characterization of IAV infection in primary human AEOs

Next, we used published methods^15,46-49^ to generate primary AEOs from human lung. We isolated human AT2 cells via FACS sorting (EPCAM^+^, Lysotracker^+^, Live cells) from otherwise healthy lung (Figure 7A, see Methods for donor details). Isolated AT2 cells were cultured in similar conditions to murine cells and using MRC5 cells as mesenchymal feeder cells. These human cultures grew faster than the murine-derived AEOs and generated organoids comprised of numerous AT2 cells (Figure 7B) without the AT1 differentiation or internal complexity or cavitation observed in mouse AEO cultures. We then proceeded to IAV infection in these human AEOs.

**Figure 7.**
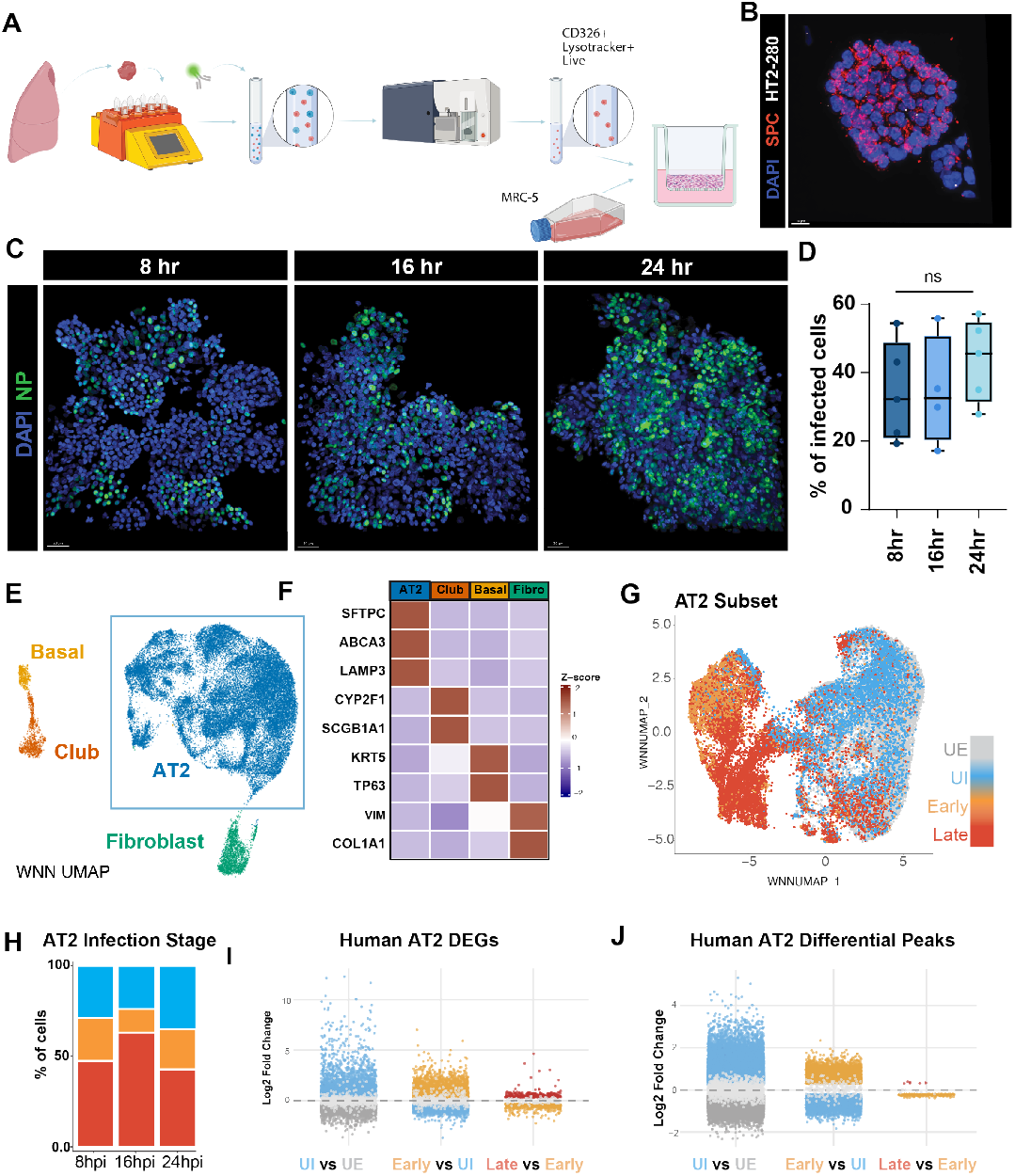
Single nuclear analysis of flu infection in Human AEOs. (A) Schematic describing the co-culture of freshly sorted human epithelial cells with MRC5 fibroblasts for creation of human AEOs. (B) IHC demonstrating clear punctate SPC expression in human AEOs; we noted relatively low level HT2-280 expression in established AEO culture. (C-D) Dynamics of PR8 IAV infection in AEOs, showing high levels of nuclear NP expression (C) which is quantified in (D). (E-F) UMAP projection (EF) and cell marker gene expression (F) of cell states found in human AEOs. (G-H) UMAP of AT2 cell subset (G) and quantification (H) showing imputed flu infection state as previously described for murine organoids. (I-J) Strip plots showing differentially expressed genes (I) or differentially accessible chromatin regions (J) in Uninfected vs Unexposed, Early Infected vs Uninfected, and Late Infected versus Early Infected human AT2 cells.

To control for viral-specific factors, we used the same PR8 influenza used for murine AEO infection. PR8 is adapted from human pandemic H1N1 but is a mouse attenuated virus and our human AEOs achieved more than double our murine average colony forming efficiency, creating dense cultures. We therefore doubled the viral load to 2.05×10^6^ PFU/well to ensure a similar MOI in human as in murine AEOs. Human organoids underwent efficient infection with more infected AT2 cells present at 8hpi and 16hpi compared to mouse AEOs, but a peak at 24h dpi similar to mouse (Figure 7C-D). We then proceeded to multiome paired snRNA and snATACseq.

Following nuclei isolation using the same protocol as for mouse AEOs, our snRNA data contained 38,610 single-cell transcriptomes with a range of 5,793 to 7,478 cells per timepoint and condition with a median of 3,527 unique molecular identifiers (UMI) (interquartile range 1,749 – 4,795) per cell, and a median of 1,927 genes (interquartile range 1,134 – 2,374) per cell. Over 75% of cells collected were identified as AT2s with the expected fibroblast presence from the MRC5 co-culture (Figure 7E-F). We observed some airway cell contamination of basal and secretory cells, which were morphologically apparent in human AEO wells and present in both control and IAV-treated cultures, suggesting they originated in the culture prior to the onset of IAV infection. Notably, we did not observe the development of an STE population in the human AEO culture. We hypothesized that the human AEO milieu was less pro-inflammatory than the murine AEO cultures; this interpretation was supported by snRNA data showing only low level upregulation of RNA for key inflammatory cytokines including TNFα, IL-1β, IL-6, IP-10/CXCL10, and RANTES/CCL5 in human AEO (Supplementary Figure S3). MRC5-derived feeder cells and the absence of an immune population resulted in expression of a smaller number of these proinflammatory cytokines, and at a lower level, than the murine primary feeder cells, leading us to conclude that the absence of STE in human AEOs may relate to an overall decrease in inflammatory signaling in non-epithelial cells following IAV infection. We can not, however, exclude the possibility that species-specific mechanisms could also be contributing to these differences in epithelial cell state.

Human AEOs contained a large population of AT2 cells within each of infection classes defined previously, so we used similar methods to classify Unexposed, Uninfected, Early, and Late Infected human AT2 cells (Figure 7G-H). Consistent with IHC data, we noted more cells classified as Late Infected even at 8hpi, with similar distribution at 16hpi and a notable decrease in both infected Late Infected populations at 24hpi, potentially suggesting a more rapid saturation of infection in human AEOs. We then performed DEG (Figure 7I) and differential chromatin accessibility analysis (Figure 7J) using the same analytical pipelines as used for the murine data. We noted similar dynamics of gene expression (Supplementary Figure S4) and chromatin accessibility in humans as in mice, with the largest changes notable comparing Unexposed to Uninfected AT2 driven by viral and immune response pathways. IAV infection led to a burst of translation-related pathways, consistent with viral replication driven by infection. This was followed by a substantial decrease in inflammatory pathways during Late infection, consistent with viral-mediated suppression as infection progresses (Supplementary Figure S4). These similar patterns implied a core conserved response to IAV across mammalian AT2 cells.

### Defining a Core, Evolutionarily Conserved Viral Response in IAV-infected AT2

To identify this core anti-viral response in AT2 after IAV infection, we developed a two-step cross-comparison of the snRNA results from the mouse and human AEOs to identify strongly conserved genes and pathways (Figure 8A). Using DEG signatures derived from pseudobulk expression profiles for each cell state (Unexposed, Uninfected, Early IAV, Late IAV) in both human and mouse, we performed pathway and GSEA analysis to identify conserved orthologues in the gene expression data (see Methods), and displayed GO pathway enrichment (Figure 8A and Supplementary Figure S5) that defined the AT2 response to IAV. We used stringent thresholds to avoid overinterpretation of sparse data and define a relatively small core conserved gene list for each stage of infection.

**Figure 8.**
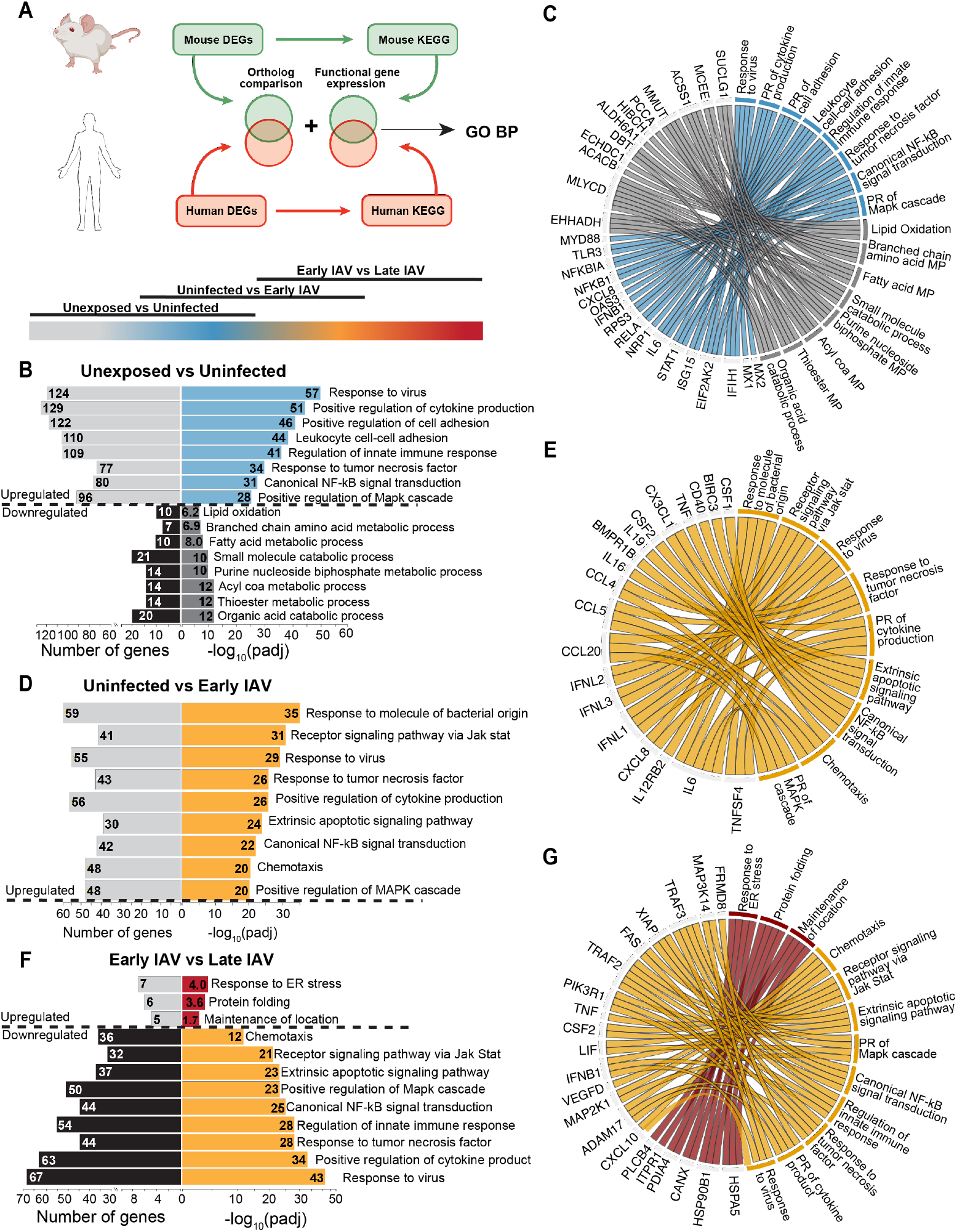
A Conserved Signature of IAV infection in AEOs. (A) Schematic depicting data analysis pipeline (see Results and Methods) used to identify core conserved regulated pathways and gene sets in IAV-infected AEOs. (B-C) Enriched and downregulated GO Terms (B) in Unexposed vs Uninfected AT2 with leading edge genes per GO terms as in (C). (D-E) Enriched GO Terms (D) in Uninfected vs Early IAV AT2 with leading edge genes per GO term as in (E). (F-G) Enriched and downregulated GO Terms (F) in Early vs Late IAV infection in AT2 with leading edge genes per GO term as in (G).

As expected, we found an early upregulation of genes associated with anti-viral and pro-inflammatory signaling in Uninfected AT2, corresponding to the primed chromatin accessibility seen in this state in both species. GO-BP output defined key pathways in viral response and immune activation (Figure 8B), with leading edge genes that defined these signatures, including NFKB, TLR3, MYD88, STAT1, and ISG15 (Figure 8C). Uninfected AT2 cells downregulated metabolic pathways necessary for surfactant metabolism, including fatty acid β-oxidation and lipid catabolic processes (Figure 8B) with these signatures defined by a decrease in critical metabolic enzymes, including ACACB and PCCA.

Comparing Early infection to Uninfected AT2, we noted no downregulated pathways and found that 87% of pathways enriched in Early infection were present in the Uninfected signature but further upregulated following infection initiation (Figure 8D-E). The remaining 13% of pathways and leading edge genes included those associated with viral infection, with an increase in vesicle trafficking, viral assembly and budding, as well as regulation of cell death. At the Late IAV stage, we noted upregulation of ER stress and protein folding response pathways, with leading edge genes including chaperones HSPA5 and HSP90B1 and the critical ER regulator CANX. Antiviral pathways are broadly downregulated, suggesting conservation of viral-mediated immune suppression in late infection across both murine and human AT2. These trends are defined by decreased expression of previously upregulated factors from the primed state, including NFKB, MAPK, IFN, apoptotic signaling, and other immune cell mediators (Fig 8F-G).

## Discussion

Here, we present data from temporal modelling of IAV infection in murine and human AEOs. These data provide both a new resource of transcriptomic and epigenomic mechanisms of IAV pathogenesis in AT2 and a technical advancement with defined protocols for viral infection and analysis in primary lung organoids. Organoid technology has enabled significant advancement in regeneration and genetic disease. To date, however, most infection in lung organoids^22,48,49^ has been primarily proof of principle, designed to verify cellular state in a new organoid method rather than disease modelling. Our data move beyond proof of principle to define a core signature of infected AT2, including loss of surfactant processing^50^ and an active response to inflammatory milieu prior to infection to prime for anti-viral response in early infection^24^. We also define a distinct shift in cellular metabolism from lipid and amino acid metabolism towards glycolysis and carbohydrate metabolism^51,52^, a metabolic change critical for viral pathogenesis^53^. Each of these characteristics mirror an important aspect of IAV-driven ARDS pathology, and demonstrate that the impact of IAV in the alveolus is not solely a product of infected cells, but also an epigenomic shift in bystander AT2s presumably in an attempt to survive in the infected milieu. When these early responses fail, IAV infection leads to hijacking of host machinery, which is recruited to translate viral proteins, resulting in increased ER stress and an upregulation of ribosomal pathways while immune signaling is suppressed. These data define key targets associated with each aspect of IAV viral pathogenesis in AT2 cells, and demonstrate that critical aspects of IAV pathogenesis and host response are conserved in AEOs.

Our data also demonstrates the capacity for uninfected AT2s to lose their AT2 cell identity and transition into a damage-associated STE state similar to previously established models of stressed and transitional lineages in the lung. Notably, these cells were detected in our murine model but not our human AEOs, and appear to be a product of the inflammatory milieu rather than direct infection. This interpretation is supported by the absence of markers of IAV infection, including Isg15 and Oas1^54^, and the absence of viral transcripts in STE in murine AEOs. Human AT2s are known to enter into STE-like states following diverse injuries, so it is of interest that they do not do so in our human AEOs. We observed a less proinflammatory milieu in human AEOs, which we speculate may be due to 1) differential fibroblast activation in primary murine mesenchyme compared to MRC5 fetal human cells, 2) absence of an immune population and reduced cytokine production in human AEOs compared to murine, or 3) a young human lung donor who may have increased resilience to viral injury at the cellular level. A limitation of our study is the presence of a single human donor for human AEOs, but multiple biological replicates and pooled temporal sequencing results resulted in a large pool of AT2 cells for study with a controlled genetic background. A key goal of the combinatorial analysis of human and mouse was to use the human signature to define conserved aspects of the more robust murine organoid modelling, allowing us to maximize the impact of the data resource without overinterpreting data from a single human donor. Our methodological advances may provide a lower barrier of entry for future studies focused on understanding the possible diversity of IAV responses in AT2 cells present across the human population.

A provocative aspect of our study is the significant epigenomic consequences for AT2 cells within an IAV-induced inflammatory milieu. Prior data had implied a key role for flu-specific gene expression in driving epigenomic changes in infected cells, but we were surprised to find the most significant changes to chromatin accessibility occurring due to inflammatory signaling in uninfected cells. Our data suggest this change is driven by immune response signals from the milieu, and is conserved in both mouse and human AEOs exposed to IAV. We speculate that this rapid epigenomic priming functions to prepare cells for a robust antiviral response following infection and is stimulated by the immune milieu when virus is detected by the host. This progression is conserved in mouse and human AT2s, implying potential therapeutic benefits for the host. Our data are also consistent with a model of epigenomic dysregulation in infected cells, notably the apparently random chromatin opening observed in early IAV infection, which may result from readthrough transcription^17^. While some IAV-infected pulmonary cells survive infection and persist in regenerating tissue^55,56^, AT2 which survive influenza B infection have reduced regenerative capacity^56^ and many infected cells are cleared by the immune system or die following viral-mediated cell lysis. Our data suggest that targeting the response of bystander cells may provide an orthogonal approach to improving alveolar resilience during viral pneumonia.

Further, it is tempting to speculate that this major impact on bystander AT2 function may underlie part of the limited impact of existing IAV treatments including neuraminidase inhibitors (NAIs) including Tamiflu (oseltamavir). Although effective at reducing symptoms when given within the first 48 hours following clinical presentation, NAIs are less effective when given later^17^ or administered in cases of severe infection^57^. This decreased effectiveness later in disease is poorly understand, and our data implies that some aspects of viral pneumonia and ARDS may have occurred due to the immune response in uninfected AT2s. It is possible that suppression of infected cells may be insufficient for symptomatic benefit after a certain period of immune activation. Given that IAV-induced ARDS remains a global clinical challenge with few effective interventions, the cellular resolution provided in our studies may provide a path forward to new therapies. The AEO model system is translatable and is highly tunable for applications such as screening for IAV therapeutics or genetic modifiers of AT2 response. Although there is still much to be learned about IAV-host interactions, future studies in primary organoid models can provide flexible, reductionist systems to identify mechanisms underlying the cellular and physiological disruptions induced by viral pathogens which drive progression to ARDS.

## Acknowledgements

The human studies in the manuscript were enabled by the gift of a pediatric lung explant, and we thank the patient and her family for their generosity in supporting pulmonary research at CCHMC. The authors would like to thank the CCHMC Single Cell Genomics Facility (especially Kelly Rangel), the Bio-Imaging and Analysis Facility (especially director Matt Kofron), and the Research Flow Cytometry Core of Cincinnati Children’s Research Foundation for extensive technical support. A special thank you to Abigail Solstad for running our PR8 plaque assay and Jordan Dale for external manuscript feedback.

## Author Contributions

*Conceptualization* – AE, WJZ

*Data acquisition* – AE, SF, HWN, BZ, AT

*Data analysis* – AE, BZ, ALZ, WJZ

*Supervision* - WJZ

*Writing – original draft* – AE, WJZ

*Writing – review and editing* - all authors

## Data and materials availability

All mouse and human organoid RNA and ATAC sequencing data has been deposited at GEO and is undergoing processing for exploration at on the NIH LungMAP portal at http://www.lungmap.net. Enrichment data used to generate all figures is available in a series of Supplemental Tables provided as a single file in the Data Supplement.

## Materials and Methods

Many of the methods used for organoid preparation and staining in this manuscript have been previously described^22^.

### Mouse ethical compliance and animals

All animal studies were conducted under the guidance and supervision of the Cincinnati Children’s Hospital Medical Center (CCHMC) Institutional Animal Care and Use Committee (IACUC) in accordance with CCHMC regulatory protocols. Mouse lines used included: C57BL/6J mice (Jackson Strain #000664), Axin2^creERT2-TdT^ (a gift from Edward Morrisey)^15,58^, and R26R^EYFP^ (B6.129×1-Gt(ROSA)26Sor^tm1(EYFP)Cos^/J; Jackson Strain #006148)^59^. All experiments included both male and female mice.

### Human ethical compliance

De-identified postmortem primary human lung tissue was obtained from the local Cincinnati organ procurement organization (LifeCenter) in accordance with UCCOM/CCHMC regulatory guidelines. While this research is not considered human studies research based on the 2018 NIH Common Revised Rule definition of Human Studies Research, tissue was accepted under UCCOM IRB# 2013-8157 (for which WJZ is a registered investigator) following approved protocols. Tissue for this study was donated by the family of a single six year-old female donor with a head injury which was not used for clinical transplantation due to a transient drop in oxygen saturation immediately prior to collection.

### Mouse PR8(Cre) Infection

Mixed background mice containing the R26R^EYFP^ were weighed and administered 0.4HAU/g of a PR8 H1N1 influenza containing Cre recombinase^55,56^ (a gift from Nicholas Heaton and Emily Hemann) at 8-12 weeks of age. After 48, 72, 96 hours, mice were weighed again and lung tissue was harvested as below.

### Mouse lung harvest for single cell suspension

Mice were anesthetized via Isoflurane exposure, followed by euthanasia via cervical dislocation and thoracotomy. The chest cavity was opened to expose the heart and lungs. The right ventricle was perfused with 5–10 mL of cold PBS to clear blood from the lungs. The perfused lungs were removed, and the individual lobes were isolated, removing as much airway as possible, and placed in cold PBS on ice. Lobes were finely chopped and transferred to a GentleMACS C tube (Miltenyi Biotec, 130-093-237) containing 5 mL of digestion buffer [PBS with Dispase, DNase, and Collagenase Type 1 as previously described]. C tubes were placed on a gentleMACS Octo Dissociator with Heaters and the following protocols were run: “*m_lung_01_02*” (36 s) twice, “*37C_m_LIDK_1*” (36 min 12 s) once, and “*m_lung_01_02*” (36 s) once. Samples were passed through a 70 µm filter (Greiner Bio-One, 542070) and centrifuged at 800 × *g* for 5 min at 4 °C. Following removal of the supernatant, 5 mL of RBC Lysis Buffer (Invitrogen, 00-4333-57) was added and incubated for 5 min. This single cell suspension was then used for flow cytometry or fluorescence-activate cell sorting as below.

### Flow cytometry of mouse lungs

Single-cell suspensions were obtained as above, and cells were resuspended in 5 mL MACS Buffer (autoMACS Rinsing Solution [Miltenyi Biotec, 130-091-222] with MACS BSA Stock Solution [Miltenyi Biotec, 130-091-376]) and passed through a 40 µm filter (Greiner Bio-One, 542040). Cells were centrifuged, the supernatant was removed, and the cell pellet was resuspended in Fc Receptor Binding Inhibitor Polyclonal Antibody (Invitrogen, 14-9161-73) diluted 1:100 in MACS buffer and incubated for 10 min at room temperature. Following centrifugation, cells were resuspended in 100uL antibody cocktail consisting of Podoplanin, BV421; CD31 eFluor450; MHC-II, BV480; CD326/EPCAM, BV785; CD24, PE-Cy7; ItgB4/CD104, AF647; CD45, AF700; Fixable Viability Dye eFluor 780; and SNA-FITC. Single color controls were generated in 1.5mL Eppendorf tubes using control cells aliquoted from vehicle/uninfected mice or, in the case of the Pr8 Cre experiment, from flu-infected mice prior to staining. Cells and controls were washed 1x with 1mL (controls) or 1-5mL (samples) of MACS buffer and the cell pellet was resuspended in MACS buffer (volume adjusted for cell count) and passed through a 35 µm filter lid (Corning, 352235) into a FACS tube. The Cytek Aurora was used for the data collection and spectral unmixing while FloJo was used for subsequent analysis.

### Murine epithelial processing for organoids

Single-cell suspensions were obtained as above, and cells were resuspended in 5 mL MACS Buffer in a mixture of the following antibodies diluted 1:100 in MACS buffer and incubated for 10 min protected from light: CD31 eFluor450, CD45 eFluor450), CD326 APC, Fixable Viability Dye eFluor 780. Cells were washed 1x with 5-10 mL of MACS buffer and the cell pellet was resuspended in MACS buffer (volume adjusted for cell count) and passed through a 35 µm filter lid (Corning, 352235) into a FACS tube for sorting. Using single-stain controls from experimental animals and wild-type littermates (TdTomato^−^) for compensation and adjusting gating to remove debris/doublets, the live/CD31^−^/CD45^−^/CD326^+^(EpCAM^+^)/TdTomato^+^ (Wnt-active AT2) population was sorted into a 1.5 mL eppendorf tube containing ‘spiked’ SAGM organoid medium.

### Fibroblast stock preparation and maintenance

Primary lung fibroblast stocks were generated from P28 C57BL/6 mice as previously described. For the use of frozen fibroblast stocks in organoids, 2-4 days prior to use in organoids, cells were thawed and replated in 10cm plates in DMEM/F12 with 10% FBS. On the day of organoid culture, fibroblasts washed, trypsinized, resuspended, and counted.

### Organoid growth medium

To generate ‘spiked’ SAGM medium for mouse lung alveolar organoids, SABM Small Airway Epithelial Cell Growth Basal Medium (Lonza, CC-3119) was combined with the following additives: SAGM Small Airway Epithelial Cell Growth Medium SingleQuots Supplements and Growth Factors (using only the BPE [2 mL], Insulin [0.5 mL], Retinoic Acid [0.5 mL], Transferrin [0.5 mL], and hEGF [0.5 mL] aliquots) (Lonza, CC-4124), Heat Inactivated Fetal Bovine Serum (Corning, 35-011-CV, final concentration 5%), Antibiotic-Antimycotic (Gibco, 15240-062, final concentration 1x), Cholera Toxin from *Vibrio cholerae* (Sigma, C8052, final concentration 25 ng/mL). For human cultures, to ‘spiked’ SAGM, T3 and Hydrocortisone additives [0.5mL ea] were added.

### Murine alveolar organoid plating and maintenance

Epithelial and fibroblasts were generated as above, and cultured in SAGM media (above) in 50% matrigel/50% SAGM on a 0.4µm Corning Transwell insert in a 24-well plate with media changes every 48h as previously described.

### Infection medium

To generate infection medium for mouse lung alveolar organoids, SABM Small Airway Epithelial Cell Growth Basal Medium (Lonza, CC-3119) was combined with the following additives: SAGM Small Airway Epithelial Cell Growth Medium SingleQuots Supplements and Growth Factors described as above for the ‘spiked’ SAGM. L-1-Tosylamide-2-phenylethyl chloromethyl ketone (TPCK)-treated trypsin from bovine pancreas (Sigma T1426) was added to a final concentration of 2ug/mL.

### Organoid Infection

Organoids for flu and controls were resuspended in 300ul/well ‘spiked’ SAGM in separate 15mL conicals if pooled or in individual Eppendorf (both techniques were used but no downstream differences in infection were detected due to the saturating MOI). At this point BSA precoating of pipette tips was not needed due to the presence of serum in the media. 1.025 × 10^6^ PFU PR8 Influenza (Charles River) was added per 300ul SAGM or per well of organoids. Organoids were then moved to an Ultra Low Adhesion Plate for incubation at 37C. At the desired timepoint, 1mL of 4% PFA was added to each well and the plate incubated at 4 °C for 45 min. Organoids of like condition were pooled in 15mL conicals and spun down at 100×*g* for 5 min at 4 °C before proceeding to permeabilization.

### Human AT2 isolation

Distal lung tissue grossly assessed as healthy for culture was isolated to avoid airway and dissected into 0.3g portions in 10cm dishes containing PBS with 1x Anti-Anti. As with the mouse lung, these portions were placed into a GentleMACS C tube and cut into small pieces before adding 5mL Digestion Buffer. Samples were digested as above and passed through a pre-rinsed 100 µm filter (Greiner Bio-One, 542000). The samples were centrifuged at 800 × *g* for 5 min at 4 °C and resuspended in 5mL RBC lysis buffer. This suspension was then filtered through a pre-rinsed 40 μm filter and spun at 500 × g for 5 min at 4 °C. The supernatant was removed, and the cells are resuspended in 500 μL of 1:10,000 FITC Lysotracker (Cell Signaling Technology 8783S) in MACS Buffer for 30min at room temperature. In the last 10 minutes, 5ul Fc receptor block in MACS Buffer was added to the suspension. 10mL MACS buffer is added and the cells spun at 500 × *g* for 5 min at 4 °C. Cells were resuspended in an antibody cocktail including EPCAM (Invitrogen 12-9326-42) and Fixable Viability Dye eFluor 780 for 10minutes on ice. Single stain controls were generated from unstained cells aliquoted before and after Lysotracker staining and before the antibody cocktail. After 1x 10mL rinse, the cell pellet was resuspended in MACS buffer (volume adjusted for cell count) and passed through a 35 µm filter lid into a FACS tube for sorting. The Live/CD31^−^/CD45^−^/CD326^+^(EpCAM^+^)/Lysotracker^+^ (AT2) population was sorted into a tube containing ‘spiked’ SAGM organoid medium for human cells (see above) at 4 °C, using a BD FACSAria Fusion cell sorter with a 100 µm nozzle.

### Human organoid culture

Human AT2s were plated as described above for murine AT2 except MRC5s (ATCC CCL-171, tested negative for mycobacterial contamination) at P8 were used as a replacement for the murine fibroblast stock. Human AT2s were then plated as described at 1:10 with P8 MRC5s in 50% Matrigel.

### Isolation of organoids for infection or whole-mount immunofluorescence

Transwells were washed (above and below) with 500 µL of ice cold PBS. Then, 500 µL of Cell Recovery Solution (Corning, 354253) was added to each Transwell, and a wide-bore pipette tip was used to mechanically disrupt the Matrigel—the mixture was pipetted up and down and transferred to a new Ultra-Low Adhesion 24-well plate (Corning 3473). Organoids were washed and isolated as previously described.

After isolation, the organoid pellet was resuspended in 1 mL of 4% PFA and incubated at 4 °C for 45 min. Permeabilization was performed with 0.1% PBS-Tween overnight at 4 °C, and blocked with 500 µL of 5% Normal Donkey Serum (Jackson ImmunoResearch, 017-000-121) in 0.1% PBS-Triton X-100 at room temperature on an orbital shaker for 1 hour (when staining with SNA-FITC, blocking was done per manufacturer instructions using Carbo-Free Blocking Solution, 10x Concentrate – Vector Laboratories SP-5040). Isolated organoids then underwent IHC as previously described using the antibodies in Supplemental Table 1 followed by clearing in fructose-glycerol clearing solution (60% vol/vol glycerol + 2.5 M fructose) and mounting. Slides were imaged immediately or stored at 4 °C.

### ELISA Analysis

Immediately prior to liberation of organoids for nuclei isolation, 50-100ul of suspension media was collected from organoid samples. These samples were analyzed with a custom Luminex panel in conjunction with the Research Flow Cytometry Facility at CCMC following the manufacturer’s protocol.

### Organoid dissociation and preparation of single-cell suspension for scRNAseq and scATACseq

Transwells were washed (above and below) with 1 mL of PBS. Then, 60 µL of organoid digest buffer (Dispase [Corning, 354235, undiluted, 50 U/mL], DNase I [GoldBio, D-301, final concentration 5 U/mL], Collagenase Type I [Gibco, 17100017, final concentration 4800 U/mL)] was added, and Matrigel plugs were gently disrupted. Nuclei were obtained following washing and digestion as previously described.

### Sequencing/library preparation

From each condition, a single nuclear preparation was prepared as described above, and a maximum of 16,000 nuclei were loaded into a channel of a 10x Genomics Chromium system by the Cincinnati Children’s Hospital Medical Center Single Cell Genomics Facility. Libraries for single nuclear multiome sequencing were generated following the manufacturer’s protocol. Sequencing was performed by the Cincinnati Children’s Hospital DNA Sequencing Core using Illumina reagents. Raw Sequencing data was aligned to the mouse reference genome mm10^25^ or human reference hg38^60^ with CellRanger-arc 2.0.2^61^ to generate expression count matrix files. To detect flu genes, a contig for each positive and negative PR8 IAV gene strand was added to each genome following 10x Genomics “Build a Custom Reference” instructions.

### Data QC and analysis

After alignment, ambient RNA contamination was addressed using scCDC (single-cell Contamination Detection and Correction)^28^, which identified contamination genes using a 0.4 restriction factor, quantified contamination ratios, and generated corrected count matrices. Using Seurat^62^ **(33)** RNA parameters from the multiome object were used to filter cells and to retain those with 300-3,500 detected features and <50% mitochondrial gene expression. Mitochondrial genes and MALAT1 were removed from downstream analysis. Doublet detection was performed using DoubletFinder^27^with optimized pK parameters determined through parameter sweeping, and only singlet cells were retained for analysis. For ATAC-seq data, Signac^61^ was used to filter cells based on ATAC fragment counts (1,800-100,000), TSS enrichment (>2), nucleosome signal (<1), and blacklist region overlap (<5%). Data integration was performed using Harmony^63^ to correct for batch effects across samples, followed by weighted nearest neighbor (WNN) analysis combining both RNA and ATAC modalities. The final dataset included cells from six samples across three timepoints (8h, 16h, 24h) under control and flu treatment conditions. Clustree^64^ and clusterProfiler^65^ were used for cluster identification.

RNA analysis was described in the main body of the paper. Briefly, we pseudobulked the AT2 cells by infection stage with equal distributions of each time point then used GSEA with DESeq2^66^ Wald statistics to determine DEGs. These were then filtered to include only log2FC of >0.38 and a adjusted P-value of <0.05. To analyze the ATAC data, we used an ArchR-inspired^39^ approach that took differential peak results identied by DESeq2 as input and then utilized ArchR’s exponential decay function and distance weighting approaches for gene activity scoring with gene size normalization. The top 25% of active genes from this data were used for ORA analysis, with the remaining significant genes serving as the detection universe for analysis. All GO-Biological Process analyses were done by filtering to pathways between 5 and 300 genes in size with significant results then filtered based on an adjusted P-value of <0.05 following FDR correction using Benjamini-Hochberg method. For human to mouse comparison, we performed KEGG pathway enrichment analysis using GSEA with DESeq2 Wald statistics as the ranking metric for each species, and generated enriched pathways for each of our infection state comparisons. KEGG pathways for each comparison were identified per species, and overlapping pathways between species were identified based on KEGG IDs. Genes contributing to the overlapping pathways for each comparison with the same directional enrichment were then used to create a gene list containing mouse and human genes that contributed to the same pathways and thus similar functional outcomes. Shared orthologs between each species’ DEGs were added to this core gene list, and the final conserved gene list was analyzed using GO-BP to determine a final conserved activity profile between each stage of infection in AT2 cells.

## SUPPLEMENTAL MATERIALS

**Supplementary Figure 1.**
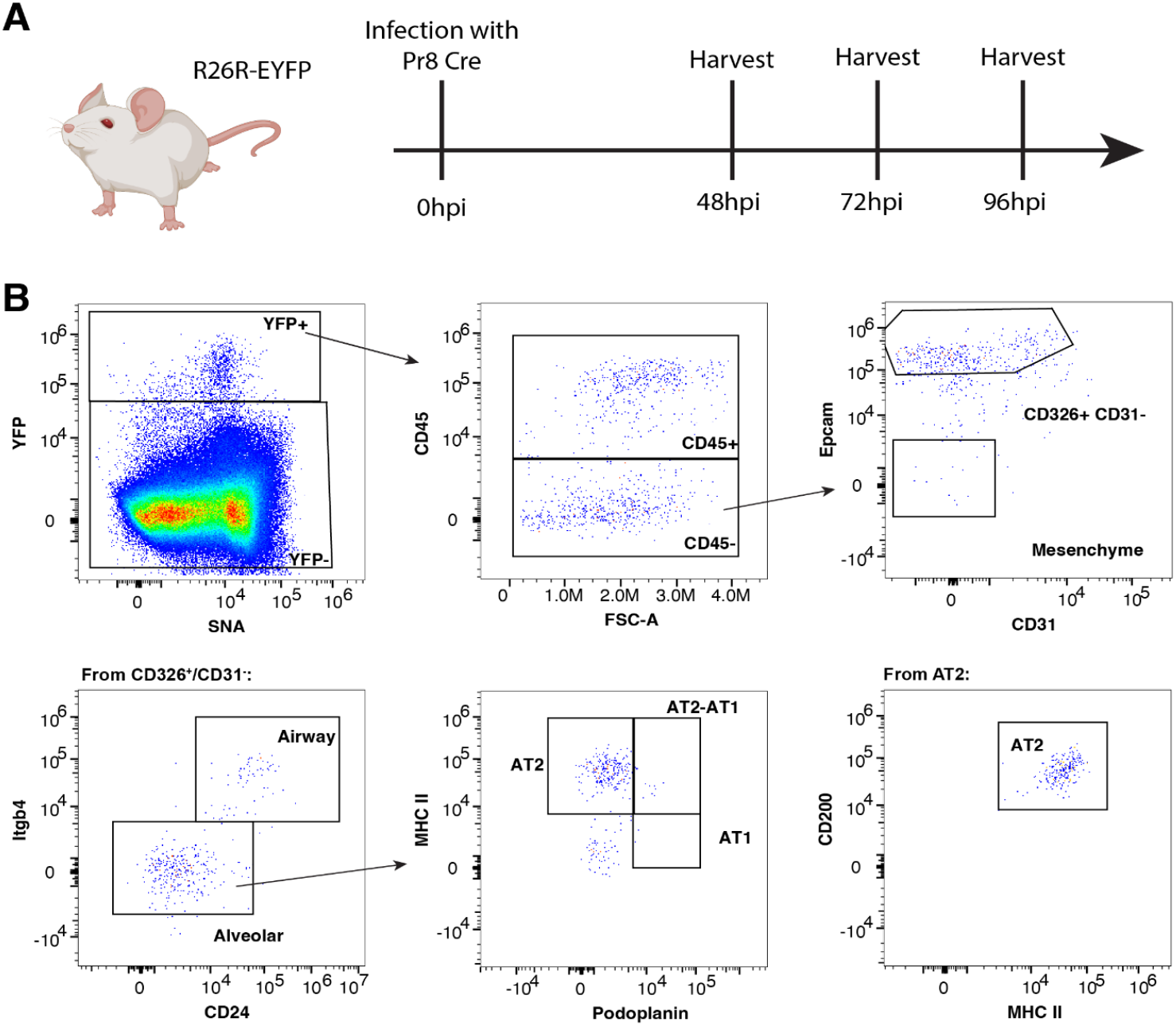
PR8-Cre infection initiation experiment and flow gating. (A) R26R^EYFP^ mice were infected with a Pr8-Cre virus and then harvested for flow cytometry 48, 72, and 96hpi. (B) Flow cytometry gating showing the identification of our YFP^+^/IAV-Infected cells with a predominance of infected AT2 cells relative to other populations

**Supplementary Figure 2.**
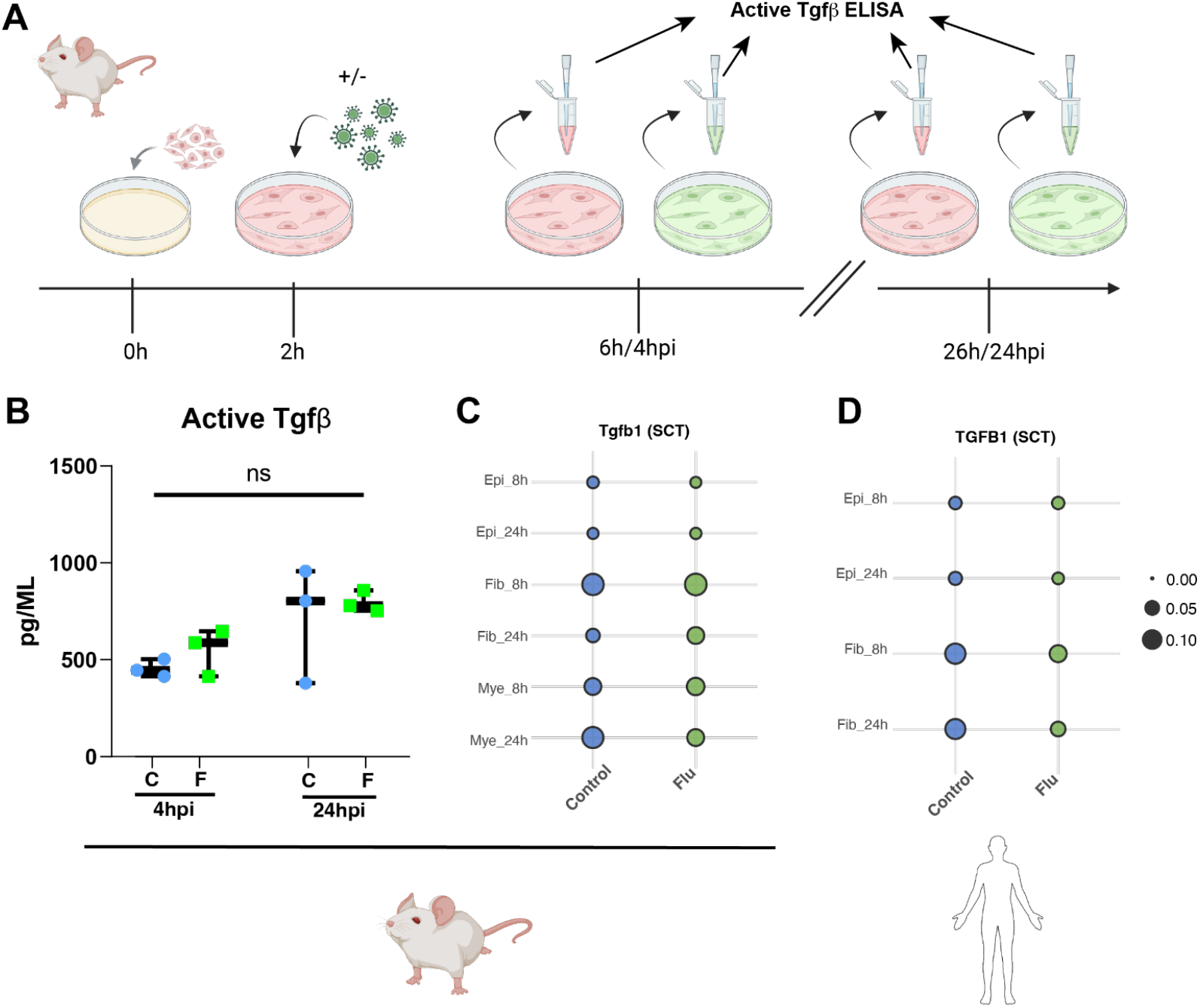
TGFβ activity and expression in the IAV-exposed milieu. (A) Schematic of D21, P3 mouse lung fibroblasts plated then cultured with or without 1.025 PFU/100k cells IAV for up to 24hpi with culture supernatant samples removed at 4 and 24hpi from IAV-exposed and control wells. (B) ELISA of culture supernatant shows an insignificant increase in active Tgfβ expression between the 4 and 24hpi timepoints and no change from control cultures. (C) Tgfb1 expression in murine AEO transcriptome data shows no difference between flu and control mean expression at 8 or 24hpi. (D) Human AEO TGFB1 expression shows no change in mean transcript expression in 8 or 24hpi.

**Supplementary Figure 3.**
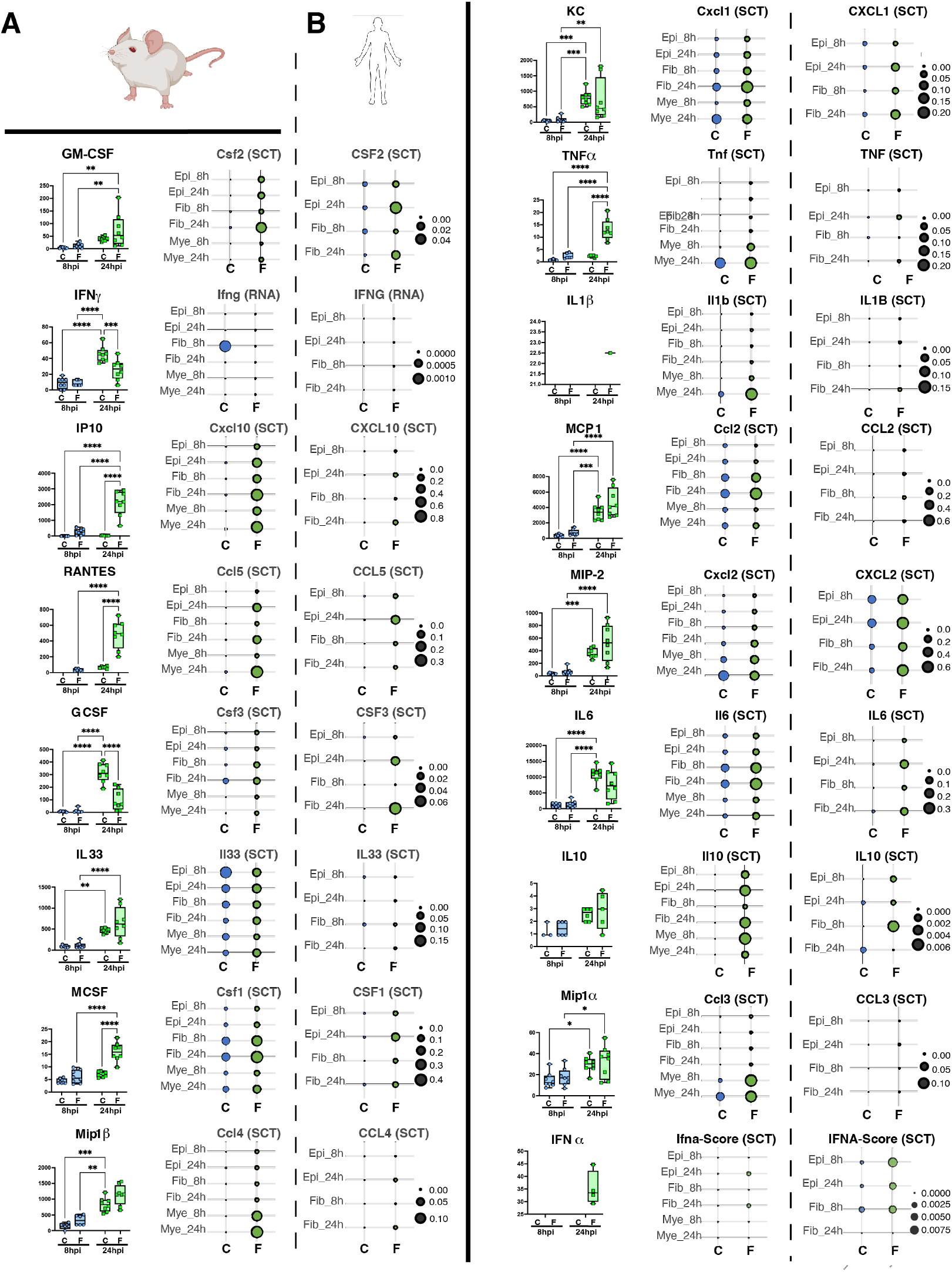
Human gene expression and Mouse ELISA and gene expression of key inflammatory cytokines. (A) ELISA data from 8 and 24hpi and unexposed AEO supernatant samples of key inflammatory cytokines, adjacent to normalized murine AEO RNA expression* within major populations (Epi = Epithelial, Fib = Fibroblast, Mye = Myeloid) reveals conserved trends between detectable protein levels and transcriptional regulation. (B) Transcription data from human AEOs indicates a unique expression signature distinct from the murine AEO culture, suggesting cellular cross-talk in the niche is critical for regulating immune signaling response. *All use assay as indicated except for Interferon alpha which was analyzed based on expression of available genes. For mice, Ifna2 and Ifna4 were detected while IFNA2 and IFNA5 were detected in the human samples for scoring. Scale for dots representing mean expression for both mouse and human data is provided to the right of each plot. P values by ANOVA with multiple comparisons: * < 0.05, ** < 0.01, *** < 0.001, **** < 0.0001. (see next page).

**Supplementary Figure 4.**
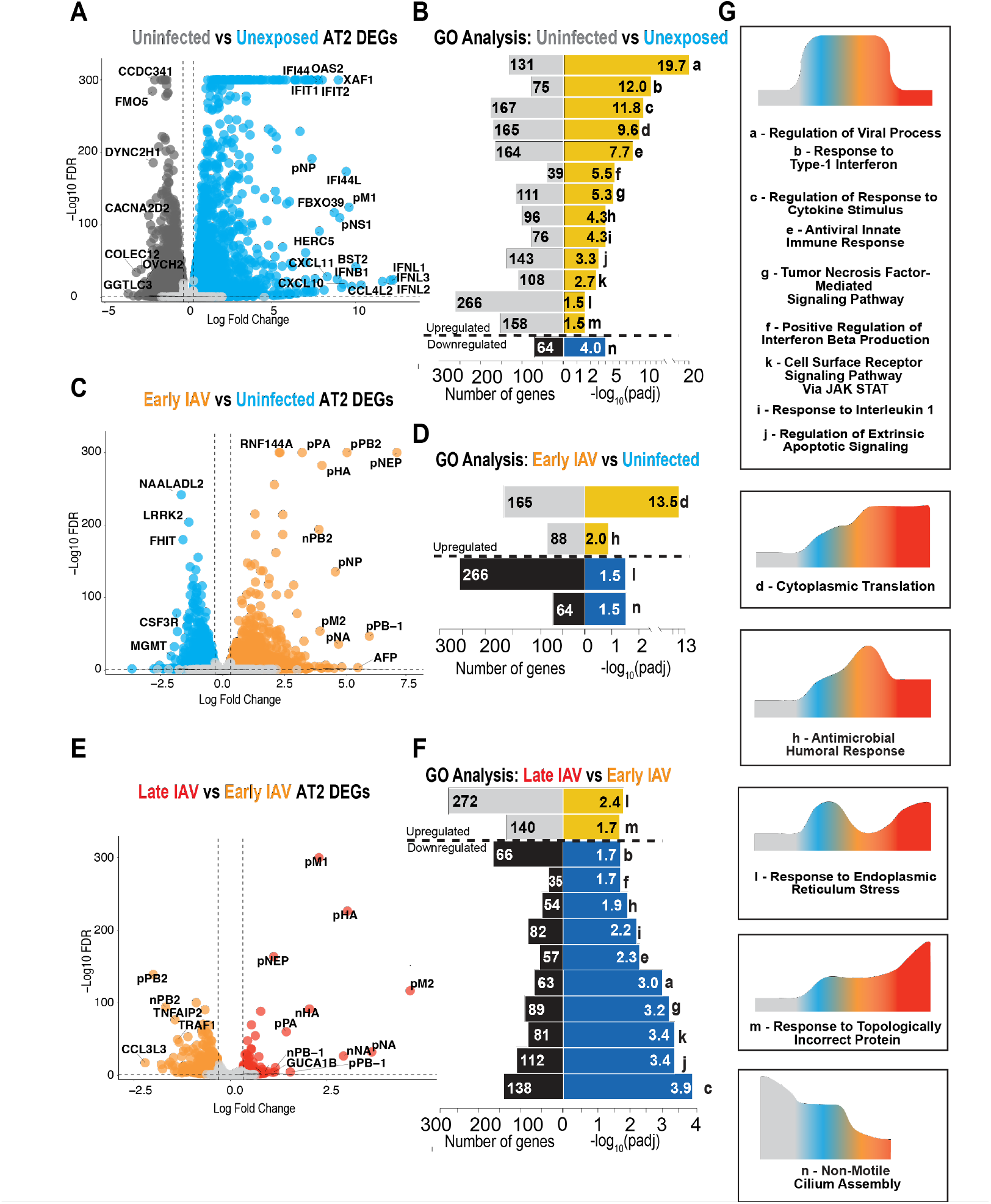
Human AT2 transcriptome analysis with GO-BP trends. (A-B) Volcano plot of DEGs (A) and GO Biological Process enrichment (B) of genes in Unexposed (grey) vs Uninfected (blue) human AT2 cells. (C-D) Volcano plot of DEGs (C) and GO Biological Process enrichment (D) of genes in Uninfected vs Early IAV (orange) human AT2 cells. (E-F) Volcano plot of DEGs (E) and GO Biological Process enrichment (F) of genes in Early IAV vs Late IAV (red) human AT2 cells. (G) Temporal dynamics of enrichment of GO terms from B, D, and F. Letter to the left of the process name matches letters to the right of the GO term bars. Some terms are enriched in multiple timepoints. Diagrams in G show the dynamics of change of associated terms (boxes) across cell states in AEOs.

**Supplementary Figure 5.**
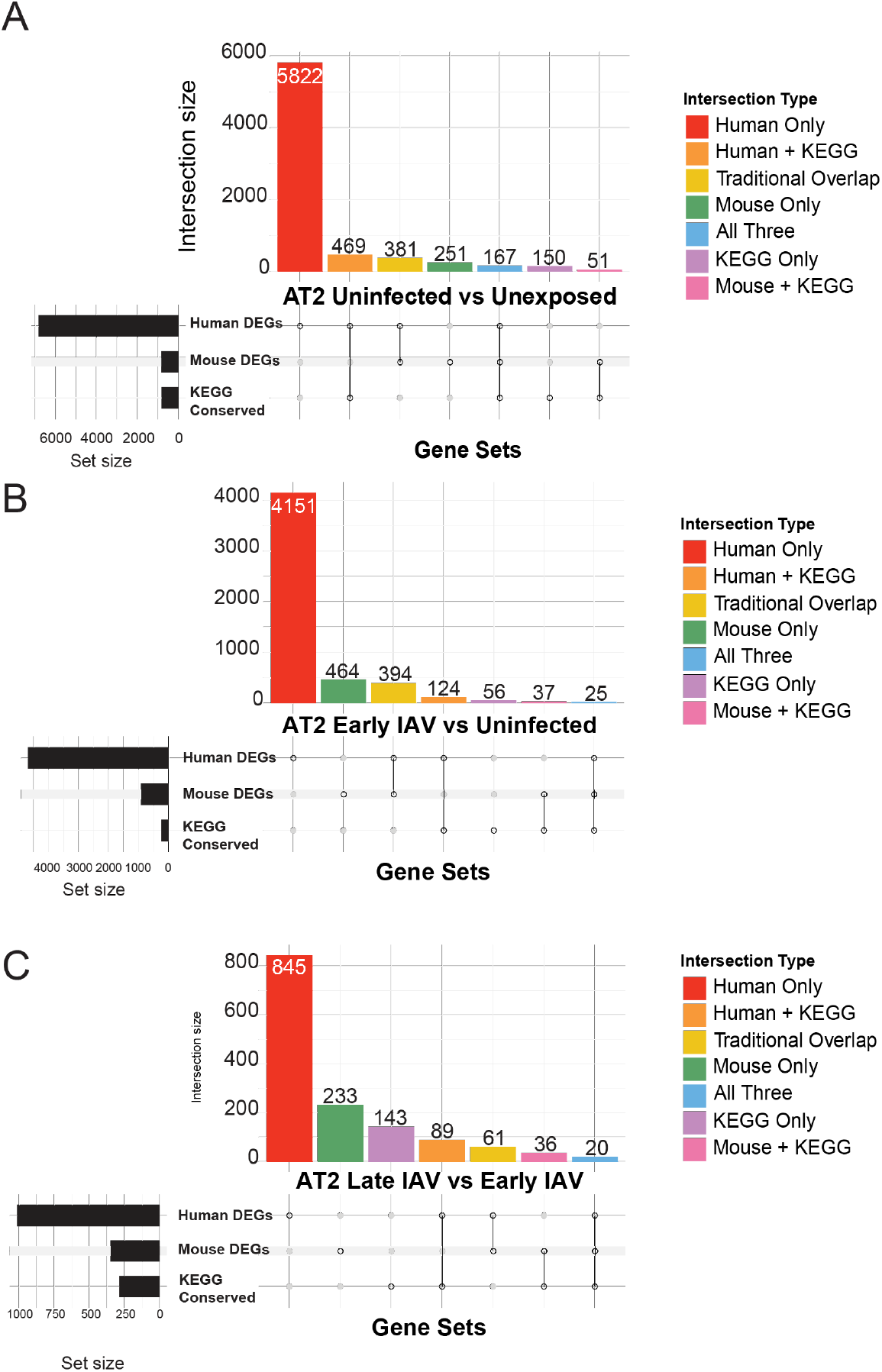
Upset plots of functional vs ortholog contribution to the murine and human integration analysis. (A-C) Upset plots of Uninfected vs Unexposed (A), Unexposed vs Early IAV (B), and Early IAV vs Late IAV (C) gene detection for mouse and human conserved IAV response analysis showing the breakdown of shared genes and pathways detected between mouse and human AT2s by comparison.

**Supplemental Table 1:** Antibodies used for organoid wholemount immunofluorescence.

| Primary antibody (1:100) | Secondary antibody (1:200) |
| --- | --- |
| **anti-RAGE** (Rat, R&D Systems, MAB1179) | **Donkey anti-Rat IgG Alexa Fluor™ 488** (Invitrogen, A21208)<br>**Donkey anti-Rat IgG Alexa Fluor™ Plus 647** (Invitrogen A48272) |
| **anti-SPC** (Guinea pig, Seven Hills, GP992 and GP993) | **Goat anti-Guinea Pig IgG Alexa Fluor™ 555** (Invitrogen, A21435) |
| **anti-Influenza NP** (Rabbit, Abcam, ab 308393) | **Donkey anti-Rabbit IgG Alexa Fluor™ 647** (Invitrogen A31573) |
| **SNA-FITC** (Invitrogen L32479) | N/A |

**Supplemental Table 2:**
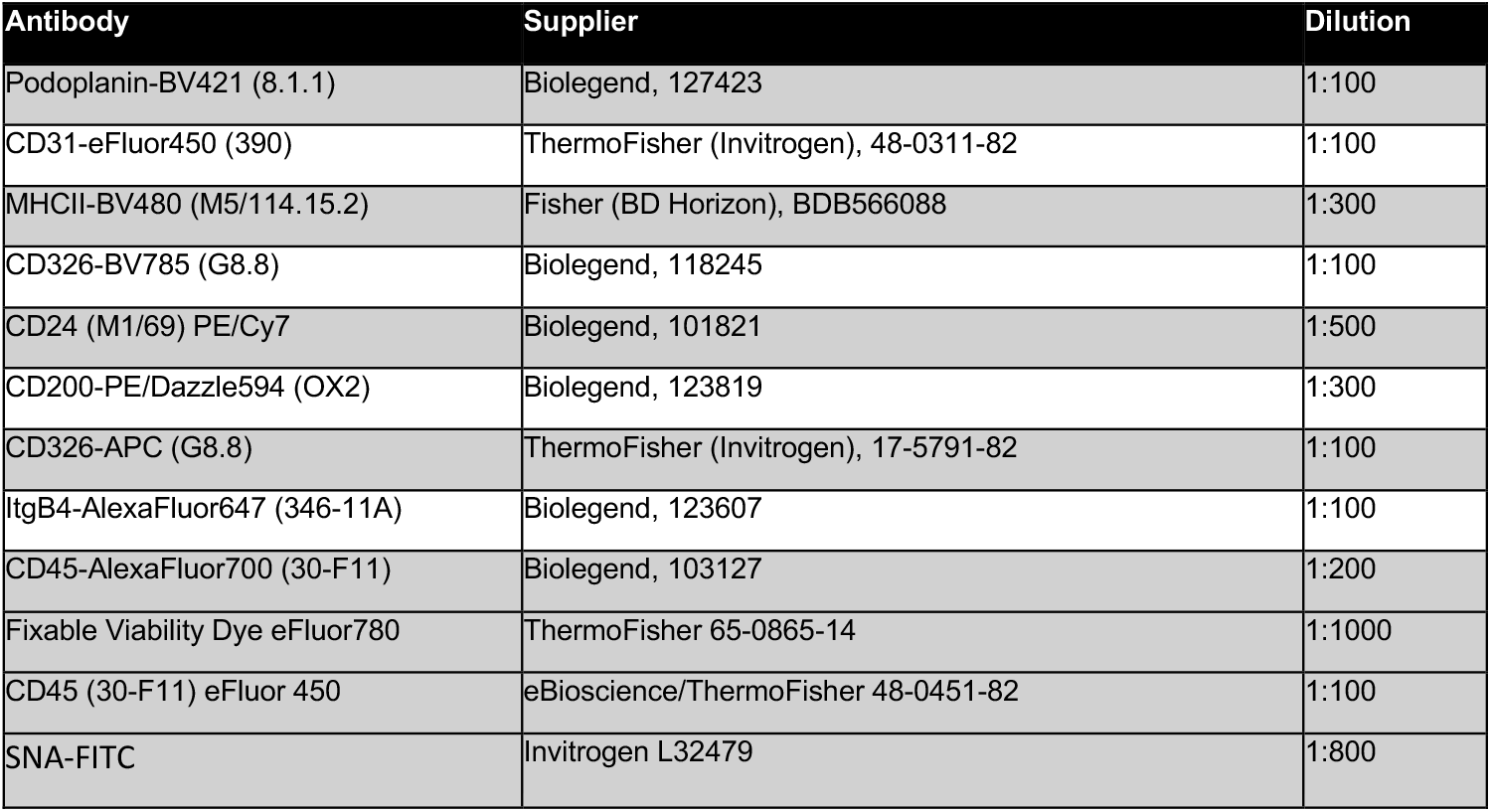
Antibodies used for FACS and spectral flow analysis.

| Antibody | Supplier | Dilution |
| --- | --- | --- |
| Podoplanin-BV421 (8.1.1) | Biolegend, 127423 | 1:100 |
| CD31-eFluor450 (390) | ThermoFisher (Invitrogen), 48-0311-82 | 1:100 |
| MHCII-BV480 (M5/114.15.2) | Fisher (BD Horizon), BDB566088 | 1:300 |
| CD326-BV785 (G8.8) | Biolegend, 118245 | 1:100 |
| CD24 (M1/69) PE/Cy7 | Biolegend, 101821 | 1:500 |
| CD200-PE/Dazzle594 (OX2) | Biolegend, 123819 | 1:300 |
| CD326-APC (G8.8) | ThermoFisher (Invitrogen), 17-5791-82 | 1:100 |
| ItgB4-AlexaFluor647 (346-11A) | Biolegend, 123607 | 1:100 |
| CD45-AlexaFluor700 (30-F11) | Biolegend, 103127 | 1:200 |
| Fixable Viability Dye eFluor780 | ThermoFisher 65-0865-14 | 1:1000 |
| CD45 (30-F11) eFluor 450 | eBioscience/ThermoFisher 48-0451-82 | 1:100 |
| SNA-FITC | Invitrogen L32479 | 1:800 |

**Supplemental File 1:** GO Term Comparisons and Leading Edge Genes For Murine and Human AEOs

